# Patient-derived response estimates from zero-passage organoids of luminal breast cancer

**DOI:** 10.1101/2024.03.24.586432

**Authors:** Róża K Przanowska, Najwa Labban, Piotr Przanowski, Russell B Hawes, Kristen A Atkins, Shayna L Showalter, Kevin A Janes

## Abstract

**Background:** Primary luminal breast cancer cells lose their identity rapidly in standard tissue culture, which is problematic for testing hormone interventions and molecular pathways specific to the luminal subtype. Breast cancer organoids are thought to retain tumor characteristics better, but long-term viability of luminal-subtype cases is a persistent challenge. Our goal was to adapt short-term organoids of luminal breast cancer for parallel testing of genetic and pharmacologic perturbations.

**Methods:** We freshly isolated patient-derived cells from luminal tumor scrapes, miniaturized the organoid format into 5 µl replicates for increased throughput, and set an endpoint of 14 days to minimize drift. Therapeutic hormone targeting was mimicked in these “zero–passage” organoids by withdrawing β-estradiol and adding 4-hydroxytamoxifen. We also examined sulforaphane as an electrophilic stress and commercial neutraceutical with reported anti-cancer properties. Downstream mechanisms were tested genetically by lentiviral transduction of two complementary sgRNAs and Cas9 stabilization for the first week of organoid culture. Transcriptional changes were measured by RT-qPCR or RNA sequencing, and organoid phenotypes were quantified by serial brightfield imaging, digital image segmentation, and regression modeling of cellular doubling times.

**Results:** We achieved >50% success in initiating luminal breast cancer organoids from tumor scrapes and maintaining them to the 14-day zero-passage endpoint. Success was mostly independent of clinical parameters, supporting general applicability of the approach. Abundance of *ESR1* and *PGR* in zero-passage organoids consistently remained within the range of patient variability at the endpoint.

However, responsiveness to hormone withdrawal and blockade was highly variable among luminal breast cancer cases tested. Combining sulforaphane with knockout of *NQO1* (a phase II antioxidant response gene and downstream effector of sulforaphane) also yielded a breadth of organoid growth phenotypes, including growth inhibition with sulforaphane, growth promotion with *NQO1* knockout, and growth antagonism when combined.

**Conclusions:** Zero-passage organoids are a rapid and scalable way to interrogate properties of luminal breast cancer cells from patient-derived material. This includes testing drug mechanisms of action in different clinical cohorts. A future goal is to relate inter-patient variability of zero-passage organoids to long-term outcomes.

## Background

The heterogeneity of luminal breast cancer [1] is difficult to capture with existing cell lines [2–4] and mouse models [4, 5] of the disease. Luminal breast cancer is partly defined by expression of the estrogen receptor (ESR1), and drugs targeting ESR1 or its ligands are a mainstay of therapy for 5–10 years after surgery [6]. For early-stage luminal cancers, it can take decades to determine whether these interventions [7, 8] are clinically effective. A faster setting for primary cancers would be highly desirable if it kept cells true to the luminal subtype.

One potential way to obtain an accelerated glimpse of cancer-cell trajectories is through patient-derived organoids [9, 10]. An early biobank of breast cancer organoids [11] preceded culture formulations geared to luminal cancers, yielding organoid lines with considerable losses in ESR1 positivity [12]. More generally, the success rate for long-term culture of luminal breast cancer organoids is only 30% [13]. Existing protocols [12] begin with whole-tumor pieces, creating a bottleneck of patient consents and non-standard handling that prevents luminal breast cancers from being fully utilized for research. Strategies for more widely establishing authentic luminal organoids would enable deeper studies of patient-to-patient variation in response to therapy or perturbations of cancer-relevant pathways.

Here, we introduce a zero-passage approach to the primary culture of luminal breast cancer organoids that retains cell identity and yields patient-specific estimates of response to molecular and genetic perturbations within two weeks of surgery. Success rates with material from primary tumor scrapes are roughly double that of previous reports [13]. The approach avoids disruptions to standardized handling of surgical specimens and miniaturizes established methods [12] to yield dozens to hundreds of organoids per replicate in six or more replicates for parallel testing. The procedure was seamlessly adopted by a 20-person team of surgeons, pathologists, and organoid culturists at the University of Virginia and could be extended to other medical research settings in the future.

## Methods

### Tissue procurement

Human sample acquisition and experimental procedures were carried out in compliance with regulations and protocols approved by the IRB-HSR at the University of Virginia in accordance with the U.S. Common Rule. In accordance with IRB Protocol #14176, scrapes of primary ESR1-positive breast tumors were collected after macrodissection with a scalpel razor. Each scrape was smeared onto a glass slide (Fisherbrand, #12-550-433) and immediately transferred into a 50 ml conical tube with 35 ml adDMEM+ (35 ml Advanced DMEM/F12 (Gibco, #12634010) supplemented with 350 μl 1M HEPES (Gibco, #15630080; final concentration = 10 mM), 350 μl GlutaMAX^TM^ (Gibco, #35050061), 350 μl penicillin–streptomycin (Gibco, #15140122; final concentration = 100 U/ml)), and 70 μl Primocin (Invivogen, #Ant-pm-1; final concentration = 100 µg/ml) and placed on wet ice.

### Plasmids

pDual_dsCas9_Venus was prepared by In Fusion cloning (Takara, #638952) using EDCPV vector (Addgene, #90085) linearized with BsmBI (NEB, #R0739S) and a PCR based insert produced by Phusion DNA Polymerase (NEB, #M0530L) containing a second sgRNA cloning site (SbfI), gRNA scaffold, and U6 promoter from pX333 vector (Addgene, #64073). Venus was replaced with mTagBFP2 (from lentiGuide-Hygro-mTagBFP2; Addgene #99374), Puromycin resistance (from pCW-Puro; Addgene, #50661; after removing its *BsmBI* by silent mutation), or Blasticidin resistance (from pCW-Cas9-Blast; Addgene, #83481) by In Fusion cloning into pDual_dsCas9_Venus digested with BamHI-HF (NEB, #R3136). Dual-sgRNAs against chromosome 1 non-coding locus, chromosome 2 non-coding locus, *NQO1*, *NFE2L2*, and *TP73* were designed with CRISPOR [14], annealed, and introduced by In Fusion cloning into pDual_dsCas9_Venus/BFP/Puro/Blast linearized with SbfI (NEB, #R0642S) or BsmBI (NEB, #R0739S). All plasmids were verified by sequencing and deposited with Addgene (#214678–214692). Oligonucleotides sequences for dual-sgRNAs, PCR cloning, and genotyping are listed in File S1.

### Organoid culture

#### Initiation

Three 50-ml conical tubes each containing two glass slides of tumor scrapes were centrifuged at 450 rcf for 5 minutes at 8°C to detach and pellet cells. The slides and supernatant were removed, and the pellet was washed with D-BSA (DMEM GlutaMAX (Gibco, #10569010) containing penicillin–streptomycin (Gibco, #15140122; final concentration = 100 U/ml) and 0.1% (wt/vol) fatty acid-free BSA (Sigma, #A6003)). Pellets were washed with D-BSA three times in 15-ml or 1.5-ml tubes depending on pellet size to remove dead cells, debris, and fat. Next, the pellet was digested in Type 1 medium [12] containing collagenase II (Fisher, #17101015; final concentration = 1 mg/ml) and ROCK inhibitor (StemCell #72304; final concentration = 10 μM) for 15 minutes at 37°C with shaking. 15-ml conical tubes were placed at a ∼20° angle in the orbital shaker at 140 rpm (New Brunswick I26 Incubator Shaker) while 1.5-ml microcentrifuge tubes were inserted in a small benchtop shaker at 350 rpm (Eppendorf Thermomixer 5350 Mixer). The digestion was stopped by adding 1/10th volume of FBS (Gibco, #16170086) directly to the tube and pipetting vigorously with a 1-ml micropipette. If large particulates were present, the sample was passed through a pre-wet 100 μm strainer (VWR, #76327102). To remove red blood cells, the cell pellet was mixed with RBC lysis buffer (Sigma, #11814389001) and incubated at room temperature for 2 min. After a final wash with D-BSA, the cells were split for organoid culture and (if possible) 2D culture. For organoid culture, the pellet was suspended in growth factor-reduced matrigel (Corning, #354230). The volume of matrigel depended on the cellular yield, with 40 μl matrigel used to resuspend a pellet of 10 μl and scaling linearly for larger pellets. The matrigel suspension was serially dispensed as one 5 μl drop per well in a 96-well plate. Paired 2D cultures were initiated by suspending cell pellets in Type 2 medium and plating on the same 96-well plate as matrigel-embedded organoids. The first 20 cases were initiated as 20 µl drops in a 48-well plate according to published recommendations [12], and volumes of 2 µl or 7 µl were tested in Fig. 4. The drops were solidified upside down in a 37°C incubator for 30 minutes and then 100 μl pre-warmed Type 2 medium [12] was added to each well.

#### Maintenance

Organoids were kept in a humidified 37°C incubator with 5% CO_2_ and refed every 2–4 of days by gently removing old media and adding fresh Type 2 medium, which was prepared weekly. For the first 20 cases when passaging was attempted,TrypLE (Gibco, #12604013) was used to dissociate the matrigel and organoids were mechanically sheared by forcefully pipetting up and down. The cells were washed with ice-cold adDMEM+, and centrifuged at 300 rcf for 5 minutes. Matrigel-free dissociated organoids were resuspended in fresh ice-cold matrigel by carefully pipetting up and down. The matrigel-suspended cells were plated in one 20 μl drop per well in a 48-well plate. The drops were solidified upside down in a 37°C incubator for 15 minutes and then 100 μl of pre-warmed Type 2 medium was added to each well.

#### Freezing and thawing

For cryopreservation, cultures were minimally digested with TrypLE to dissolve the matrigel but retain the organoids intact. Organoids were washed with ice-cold adDMEM+, and centrifuged at 300 rcf for 5 minutes. The matrigel-free dissociated organoids were resuspended in Recovery Cell Freezing Medium (Gibco, #12648010) and transferred to a –80°C cell-freezing container for at least overnight before transferring to liquid nitrogen. Organoid cryovials were thawed rapidly in a 37°C water bath, and organoids were diluted in pre-warmed adDMEM+ supplemented with ROCK inhibitor (final concentration = 10 μM). After centrifugation at 300 rcf for 5 minutes at 4°C, the supernatant was removed and the pellet of thawed organoids was plated as described above.

#### Chemical perturbation

For hormone interventions, β-estradiol was removed at day 5 from Type 2 medium and DMSO (Sigma, #D2650; final concentration = 0.1%) or 4-hydroxytamoxifen (Sigma, #H7904; final concentration = 200 nM or 3 μM where indicated) was added. The fresh Type 2 medium ± drug was exchanged on days 7, 10, and 12. For nutraceutical treatment, 10 μM sulforaphane (Sigma, #S4441) was added on day 7 with medium exchange on day 10.

#### Phenotyping

Brightfield images were scored for any evidence of plastic attachment or organoid death on each day of culture. For viability staining, organoid cultures at the 14-day endpoint were incubated with DAPI (Invitrogen, #D1306; final concentration = 0.1 μg/ml) and NucView (Sigma, #SCT100; final concentration = 5 μM) in DPBS (Gibco, #14190144) for one hour in 4°C, and fluorescence images were digitally acquired on an EVOS M7000 (ThermoFisher, AMF7000) with DiamondScope software (version 2.0.2094.0). The total counts of EGFP and/or TdTomato positive organoids were collected by digital image acquisition on an EVOS M7000 (ThermoFisher, AMF7000) with DiamondScope software (version 2.0.2094.0) and combined with segmented estimates of overall organoid size.

### Cell culture

MCF7 (ATCC, #HTB-22) were cultured in EMEM (ATCC, #30-2003) supplemented with 10% FBS, 1% penicillin–streptomycin, and 10 µg/ml insulin (Gibco, #12585014). T47D (ATCC, #HTB-133) were cultured in RPMI-1640 (ATCC, #30-2001) supplemented with 10% FBS, 1% penicillin– streptomycin, and 0.2 Units/ml insulin. CAMA1 (ATCC, #HTB-21) were cultured in EMEM supplemented with 10% FBS and 1% penicillin–streptomycin. EFM19 (DSMZ, #ACC231) were cultured in RPMI-1640 supplemented with 10% heat-inactivated FBS and 1% penicillin–streptomycin. HCC1500 (ATCC, #CRL-2329) were cultured in RPMI-1640 supplemented with 10% FBS and 1% penicillin– streptomycin. ZR-75-1 (ATCC, #CRL-1500) were cultured in RPMI-1640 supplemented with 10% FBS and 1% penicillin–streptomycin. MCF10A-5E [15] were cultured in DMEM/F12 (Gibco, #11330-032) supplemented with 5% horse serum (Gibco, #16050122), 20 ng/ml EGF (Peprotech, #AF-100-15), 500 ng/ml hydrocortisone (Sigma, #H0888), 100 ng/ml cholera toxin (Sigma, #C8052), 10 µg/ml insulin (Sigma, #I1882), 1% penicillin–streptomycin. Cells were grown and passaged according to ATCC or DSMZ guidelines. Cell images were digitally acquired on an EVOS M7000 (ThermoFisher, AMF7000) with DiamondScope software (version 2.0.2094.0).

### Organoid growth measurement

Organoid images were digitally acquired every 2–3 days on an EVOS M7000 (ThermoFisher, AMF7000) with DiamondScope software (version 2.0.2094.0) and analyzed with OrganoSeg [16]. The image sets were segmented using the following parameters: no out of focus correction, intensity threshold = 0.5, window size = 80, and size threshold = 150 (except for UVABCO61 in Fig. 7 where size threshold = 300). Area measurements were exported as a .xls file and further analyzed as described below for each experiment.

### RNA isolation and RT-qPCR

RNA from primary tumors, organoids, 2D cultures, and cell lines was isolated with TRIzol reagent (Life Technologies, #15596018) followed by purification with the Direct-zol RNA MicroPrep Kit including DNase treatment (Zymo Research, #R2062). For organoids, matrigel was removed by TrypLE digestion before cell lysis. 25 ng of RNA was reverse transcribed with 250 ng oligo(dT) (Invitrogen, #18418012), 25 pmol random hexamer (Invitrogen, #N8080127), and 200 units of Superscript III (Invitrogen, #18080044). Specific transcripts were measured by quantitative PCR using 1.3 µl cDNA template (1.67 ng of original RNA per reaction) and 3.75 pmol each of forward and reverse primers together with a homemade master mix used at a final concentration of 10 mM Tris-HCl (pH 8.3), 50 mM KCl, 4 mM MgCl_2_, 200 µM each of dATP, dCTP, dGTP and dTTP, 150 µg/ml BSA, 5% glycerol, 0.25x SYBR green (Invitrogen, #S7563), and 0.025 U/ml Taq polymerase (NEB, #M0267) in a final reaction volume of 15 µl and published thermal cycling parameters [17]. Gene abundances were calibrated against purified amplicons and normalized to *B2M* as a housekeeping gene. Primer sequences are listed in File S1.

### RNA-seq

#### Sequencing and pre-processing

RNA-seq was performed by the Genome Sciences Laboratory at Center for Public Health Genomics with 50–550 ng total RNA (TruSeq® Stranded Total RNA Library Prep, Illumina, #20020596) on an Illumina NextSeq instrument (500/550 High Output Kit v2.5 (150 Cycles), #20024907). RNA-seq data was aligned to the human assembly GRCh38.86 and quantified with STAR v2.5 [18]. Gene counts were batch corrected with ComBat_seq [19] taking each patient as a batch.

#### Cell type deconvolution

CIBERSORTx [20] was used to infer fractions and expression profiles of different cell types in each sample. A signature matrix was built from scRNA-seq data of human breast cancers [21] analyzed with Seurat v5 [22]. After clustering, cell clusters not assigned to B cells, CD4^+^ T cells, CD8^+^ T cells, macrophages (MΦs)/ dendritic cells (DCs), adipocytes, stroma, endothelial cells, muscle cells, luminal cells, or basal cells were discarded. Data were downsampled to contain no more than 2000 cells per cell type before using CIBERSORTx to create the signature matrix and gene expression profile (GEP) file. The signature matrix was used with the bulk RNA-seq data to infer cell fractions. For differential expression analysis, the signature matrix and GEP file were used to infer mean expression and standard error for tumor scrapes, zero-passage organoids, 2D cultured cells and cell lines for grouped basal and luminal cells (these two were relabeled as “LuminalBasal” and other classes were relabeled as “NotLuminalBasal” as an optional classes file for CIBERSORTx). To calculate the proliferation index in every sample, CIBERSORTx was used in high-resolution mode with the same signature matrix, GEP file, and classes file.

#### Proliferative status

A proliferation index was calculated as in [23] with a combination of 157 genes correlated with cell proliferation rates. 32 out of 157 genes were not inferrable by CIBERSORTx and thus assigned a constant estimate in the calculation.

#### Gene set enrichment analysis

To assess hallmark pathways, gene set enrichment analysis was performed using the fgsea R package [24] on fold change ranking of differentially expressed genes between tumor scrapes and the combined basal–luminal expression inferred by CIBERSORTx in GEP mode.

### DNA isolation and genotyping

Genomic DNA for CRISPR/Cas9 genotyping was isolated from cell lines and organoids by using Quick Extract DNA Extraction Solution (Lucigen, #QE09050). For organoids, matrigel was removed by TrypLE digestion before DNA isolation. Single organoids were hand-picked with capillary pipet tips and examined for locus deletion by PCR using MyTaq™ Red Mix (Bioline, #BIO-25043) and the following cycling parameters: 95°C for 1 minute, 45 cycles of 95°C for 15 seconds, 60°C for 20 seconds, 72°C for 30 seconds, followed by 72°C for 5 minutes. Oligonucleotides sequences are listed in File S1.

### Lentiviral transduction

#### Packaging

Preparation of lentiviruses by CaPO_4_ precipitation was less efficient for pDual vectors and not compatible with alternative base media. Therefore, we used lipofection (Lipofectamine 3000, #L3000001) of HEK293T cells to package psPAX2 (Addgene, #12260), pMD2.G (Addgene, #12259), and pDual_dsCas9_Venus/BFP/Puro/Blast with different sgRNA combinations or pLX302 EGFP-V5 puro (Addgene, #141348) and pLX302 TdTomato-V5 puro (Addgene, recombined from #82404). For organoid transduction, virus was prepared in high-glucose DMEM (Gibco, #11965092) supplemented with 10% FBS and 1% penicillin–streptomycin. Virus was collected at 48 hours, passed through a 0.45 µm filter, concentrated with Lenti-X™ Concentrator (Takara, #631231), and resuspended in 1/10th volume of fresh Type 2 medium [12]. For unconcentrated transduction of cell lines, pDual lentivirus was prepared in the culture medium recommended for the transduced cell line.

#### Primary cell transduction

Breast cancer organoids were first transduced during organoid initiation after RBC lysis. Cells were incubated with 10x concentrated virus in Type 2 medium for one hour at 37°C with shaking as described for collagenase treatment before, after which the initiation protocol was continued. Second, upon matrigel solidification, 100 μl of 10x concentrated virus in Type 2 medium was added to the well (instead of unconditioned Type 2 medium) for 72 hours along with polybrene (Sigma, #H9268; final concentration = 8 µg/ml). After transduction with pDual_dsCas9_Venus, Shield-1 ligand (Aobious, #AOB1848; final concentration = 200 nM) was added at day 0, replenished during refeeding at days 3 and 5, and removed upon refeeding at day 7.

#### Cell line transduction

MCF7 (6 x 10^5^ cells/well) and T47D (5 x 10^5^ cells/well) were seeded in a 6-well plate for 24 hours and then 1x virus in the recommended culture medium was added with polybrene (8 µg/ml) and Shield-1 ligand (200 nM) for 72 hours. After transduction, pDual_dsCas9_Venus/BFP cells were sorted for Venus or BFP on an Influx Cell Sorter, and pDual_dsCas9_Puro/Blast cells were selected in puromycin (MP Biochemicals, #100552; final concentration = 2 µg/ml) or blasticidin (Gibco, #R21001; final concentration = 10 µg/ml) until control plates had cleared.

### Immunoblotting

Total protein from the cultures was extracted with radioimmunoprecipitation assay buffer (50 mM Tris-HCl (pH 7.5), 150 mM NaCl, 1% Triton X-100, 0.5% sodium deoxycholate, 0.1% sodium dodecyl sulfate, 5 mM EDTA supplemented with 10 µg/ml aprotinin, 10 µg/ml leupeptin, 1 µg/ml pepstatin, 1 mM phenylmethylsulfonyl fluoride, 1 µg/ml microcystin-LR, and 200 mM sodium orthovanadate), separated on 10% polyacrylamide gels, transferred to polyvinylidene difluoride membranes, blocked with 1x Odyssey blocking solution (LI-COR, #927-70001) for one hour, probed overnight with primary antibodies followed by IRDye-conjugated secondary antibodies for one hour, and scanned on a LI-COR Odyssey instrument with Odyssey software (version 3.0) as previously described [25]. Primary antibodies recognizing the following antigens were used at the indicated dilutions: p73 (Abcam, #ab40658; 1:500), HSP 90ɑ/β (Santa Cruz Biotechnology, sc-13119; 1:1000). Secondary antibodies were used at the indicated dilutions: goat anti-rabbit (LI-COR, #926-68071; 1:20,000) and donkey anti-mouse (LI-COR, #926-32212; 1:20,000).

### Statistical analysis

#### Proportional hazards modeling

Cox Proportional Hazards Modeling was used to investigate the association between the success of organoid culture and the available deidentified patient data. The survival R package was used for multivariate Cox regression analysis and the survminer R package was used to generate the forest plot for culture success in RStudio (version 2023.06.2) and R (version 4.2.1). If tumor grade or estimated size were missing, those values were inferred based on the mean of the rest of the samples. If ESR1 and PGR percentages were annotated as above specific percentage, 0.5% was added to the named number (for example >95% was analyzed as 95.5%). Plastic attachment and death events were analyzed by a Cox proportional-hazards model of scores collected on days 2– 14 with right censoring used to indicate samples without events on day 14. The model considered organoid source (*n* = 12 patients) and culture volume (2 µl or 5 µl) as independent categorical variables.

#### Multiway ANOVA

Cross-sectional area measurements were shifted by subtracting the size threshold used for image segmentation, adding a size offset of 10 pixels^2^, and log transforming to achieve a normal distribution. The anovan function in MATLAB (version R2023a) was used on the shifted log-transformed areas to track the fixed effects of patient, culture day, culture volume (nested within patient), and technical replicate (nested within volume) on the shifted log transformed area. Pairwise interactions between factors were included, and type 3 sum of squares assessed residuals after considering main effects and interaction terms. The volumes were analyzed together and as pairs, which were Šidák-corrected for multiple-hypothesis testing.

#### Organoid growth modeling

We began with an equation for exponential volumetric growth defined by a characteristic doubling time: 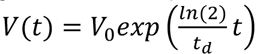, where *V*(*t*) describes the change in volume over time, *V*_0_ is the volume at *t* = 0, and *t_d_* is the doubling time in days. For a spherical organoid, volume 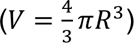 and cross-sectional area (*A* = π*R*^2^) are proportionally related: *A*(*t*) ∼ *V*(*t*)^2/3^.) Therefore: 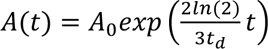, where *A*_0_ is the cross-sectional area at *t* = 0. Mean cross-sectional area (calculated after shifted log transformation) for each study was regressed against *A*(*t*) by nonlinear least-squares with the lsqcurvefit function in MATLAB. The data were fit globally to constrain a single estimate of *A_0_* across common genotypes. For the hormone-intervention study, all conditions shared the same *A_0_* because the perturbations did not begin until *t* = 5 days. For the neutraceutical study, the control, dual sgChr1, and dual sgNQO1 organoids were each fit with their own *A_0_* because transductions began at *t* = 0 days. Confidence intervals were estimated by support plane analysis, whereby either *A_0_* or *t_d_* was displaced from its best fit, the remaining parameters were re-estimated, and the decrease in model quality was assessed by *F* test of the residuals scaled by the degrees of freedom.

## Results

### Reconfigured gross processing of breast tumor resections

Breast lumpectomies are typically placed in formalin right after excision in the operating room or upon arrival to surgical pathology (Fig. 1A, solid black to orange). There are alternatives for preserving viable cells, but the amount of material requires patient consent and pre-defined eligibility criteria, which encumbers daily clinical practice. In pursuit of more flexibility, we hastened the first step by transferring fresh material to pathology within 30 minutes (Fig. 1A, dashed black). The delay is negligible with respect to immediate fixation because formalin penetrates less than 0.5 mm into the bulk tumor over this time [26]. During gross processing, cross-sections of the tumor are smeared on uncharged microscope slides before they are placed in formalin (Fig. 1A, dashed orange). The additional step does not divert any bulk material from pathology, allowing these tumor scrapes to be considered as medical waste. Tumor scrapes are compatible with most biochemical and immunocytochemical methods (Fig. 1A, blue). Importantly, they yield as many as several thousand viable carcinoma cells, which can be isolated off the slides and cultured as organoids according to protocols for tumor pieces [12, 13]. Eligibility to proceed with organoid culture is dictated by the overall cellularity of the scrape, which cannot be defined preoperatively. Therefore, our Institutional Review Board has granted this study a waiver of consent under 45CFR46.116 of the 2018 Common Rule. Together, the coordinated reordering of steps between surgeons, scientists, and pathologists enabled many fresh luminal breast cancers to be tested for organoid growth and survival.

**Figure 1.**
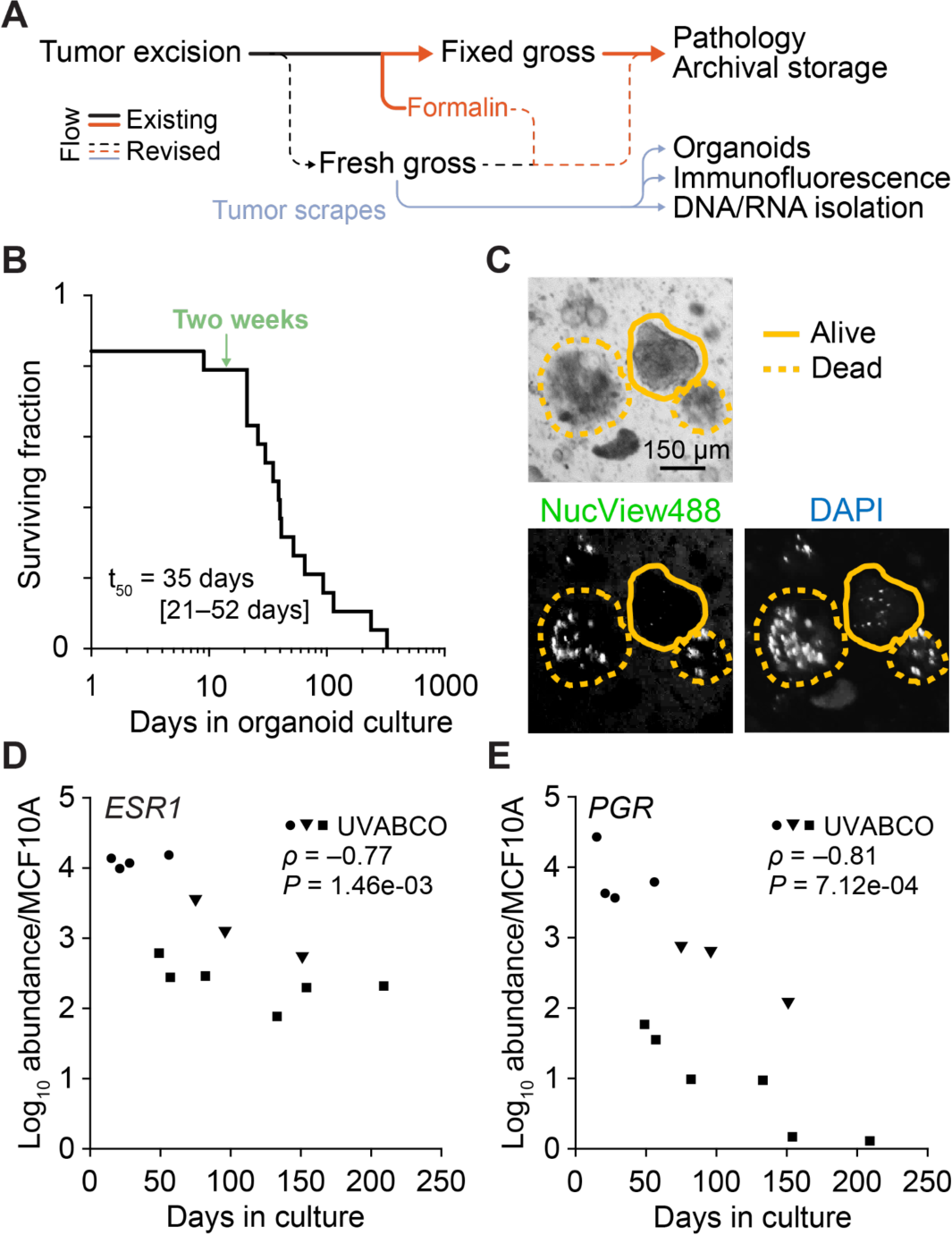
Loss of viability and hormone-receptor expression in organoids isolated from tumor scrapes. **(A)** Schematic detailing the standard gross processing of surgically resected tumors (solid black-to-orange line) alongside changes (dashed lines) that incorporate tumor scrapes (blue line). The addition of formalin is in orange. **(B)** Kaplan-Meier survival analysis of luminal breast cancer organoids generated from tumor scrapes (*n* = 19 independent luminal breast cancers). The median survival time (t_50_) is shown with the 95% confidence interval in parentheses, and the two-week time point is annotated (green). **(C)** Live-cell NucView® 488 Caspase-3 and DAPI staining of live (solid line) and apoptotic (dashed line) organoids. Scale bar is 150 µm. (**D** and **E**) RT-qPCR of *ESR1* and *PGR* expression in organoids from three patients (UVABCO92, circles; UVABCO1, squares; UVABCO5, triangles) at different passages over time. Relative transcript abundance is normalized to MCF10A-5E as a basal control.

### Prolonged culture of luminal tumor organoids causes death and loss of hormone receptors

Breast tumors of the luminal A subtype are less proliferative and much more prevalent than luminal B tumors [27]. Luminal tumors are also often wild-type for TP53 [28] and therefore remain poised to undergo apoptosis. Accordingly, despite ∼80% success in initiating luminal breast cancer organoids from tumor scrapes (*n* = 20 cases), we found that the replicative life span of primary tumors was limited. The median survival time was 35 days, and none of the cultures lasted more than one year (Fig. 1B). When terminal cultures were stained with a caspase activity reporter and a viability dye, we found that most organoids died by apoptosis (Fig. 1C). Organoid culture supports proliferation of luminal cancer cells, but the accumulating replicative stress apparently prohibits their indefinite maintenance.

A defining characteristic of luminal breast cancers is expression of the hormone receptors, ESR1 and PGR, which often disappear when luminal cells are cultured on tissue-culture plastic [29, 30]. Using three cultures that persisted for at least two months, we tracked hormone-receptor transcript abundance over multiple organoid passages. Consistent with recent work [12, 31], *ESR1* and *PGR* declined exponentially with time in culture even as cells remained viable (Fig. 1D,E). The drift may reflect de-differentiation of the luminal cancer or outcompetition by a subpopulation of hormone-negative cells [30, 32]; regardless, the results indicate that luminal tumor organoids deviate rapidly from the primary tumor.

### Zero-passage organoids preserve luminal characteristics of primary tumors

The surviving fraction of luminal cases was stable for the first three weeks (Fig. 1B) before most organoids needed to be dissociated and split to reduce cell density [12]. We thus pursued an accelerated format with a pre-designated endpoint of 14 days. The intent of this “zero-passage” approach was to capture a few culture-induced cell doublings in response to perturbations of interest while hopefully minimizing drift of the luminal state.

We tested this premise by initiating a second set of organoids (*n* = 18) and quantifying *ESR1* and *PGR* abundance relative to the originating tumor scrape (Fig. 2A,B). Six luminal breast cancer lines and one nontransformed basal line (MCF10A) were included as positive and negative controls respectively for each hormone receptor in the quantitative PCR. From six scrapes, we successfully maintained cells for 14 days as a monolayer on tissue-culture plastic with organoid culture medium. These organoid-paired samples distinguish the contribution of i) extracellular matrix ligands in matrigel, ii) the compliance of matrigel, and iii) the microenvironment effects of cell encapsulation to the molecular profiles observed. Although decreases in both hormone receptors were often measured relative to the corresponding tumor scrape, zero-passage organoids retained hormone receptor expression within the clinical range measured across patients: 21–1000-fold above MCF10A for *ESR1* and 20–8700-fold above MCF10A for *PGR* (Fig. 2A,B; yellow). Importantly, monolayer-cultured cells (2D) had significantly lower *ESR1* and *PGR* abundance compared to zero-passage organoids derived from the same tumor scrape (Fig. 2A,B; blue). Recognizing that hormone receptor abundance would continue to decline past 14 days (Fig. 1D,E), these results supported zero-passage organoids for short-term examination of most luminal breast cancers.

**Figure 2.**
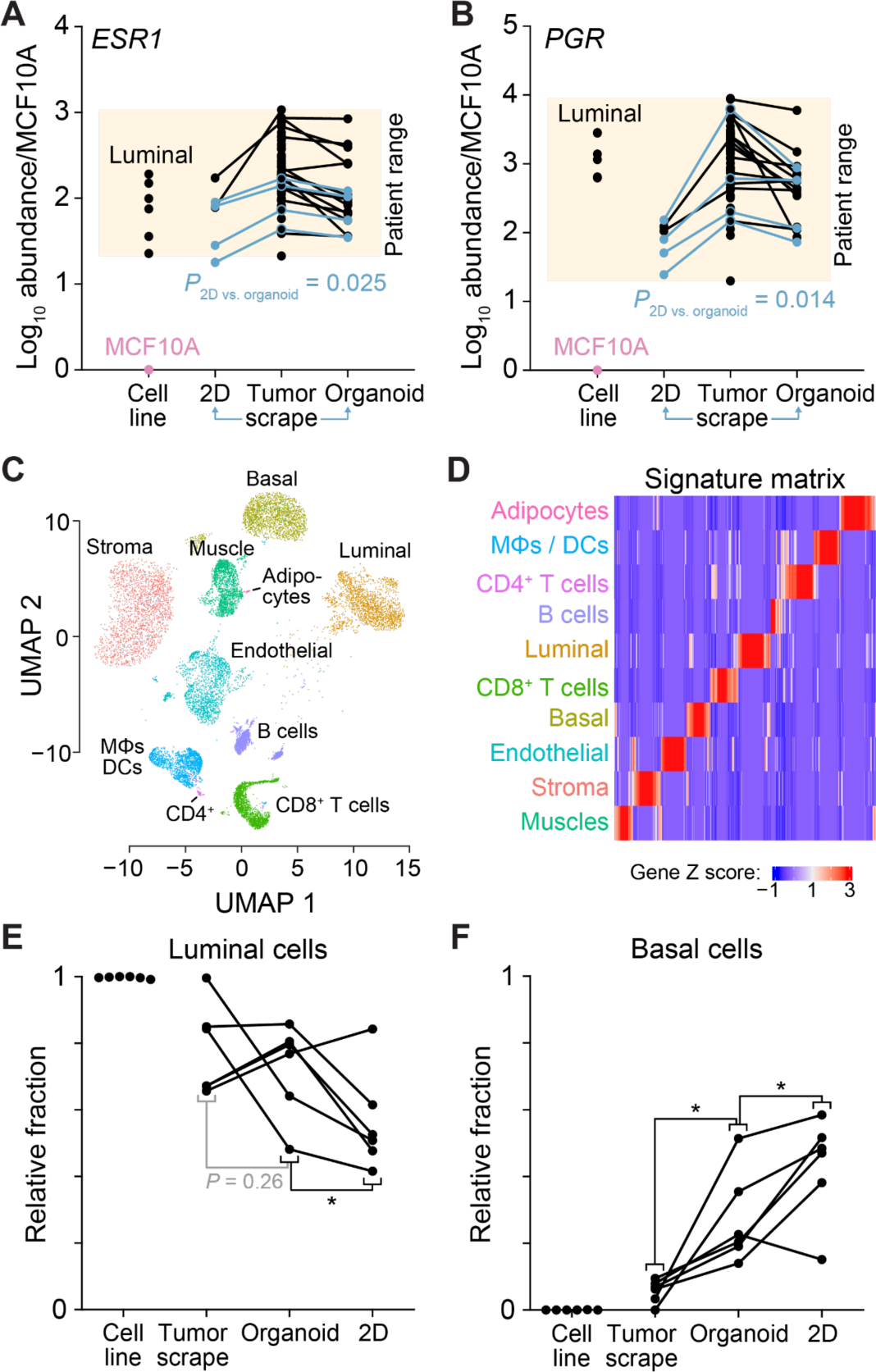
Zero-passage organoids preserve short-term hormone-receptor expression and reduce de-differentiation of luminal breast cancer cells. (**A** and **B**) Comparison of *ESR1* and *PGR* expression in originating tumor scrapes, zero-passage organoids, and 2D cultured cells relative to a basal control (MCF10A-5E; pink) and luminal breast cancer cell lines (MCF7, T47D, HCC1500, CAMA1, EFM19, and ZR-75-1). Shaded boxes indicate the observed range of expression among patients. Paired samples are connected with black lines. Blue highlights paired samples of tumor scrape, 2D culture, and zero-passage organoids (*n* = 4 independent luminal breast cancers). Paired 2D and organoid cultures were compared for *ESR1* and *PGR* abundance by paired *t* test after log transformation. **(C)** Downsampled scRNA-seq data of human breast cancers [21] used to define a CIBERSORTx signature matrix for bulk RNA-seq deconvolution [33]. **(D)** Gene-by-cell type signature matrix for deconvolving lineages in tumor scrapes, zero-passage organoids, and 2D cultures. (**E** and **F**) Comparison of luminal and basal lineage fractions in originating tumor scrapes, zero-passage organoids, and 2D cultured cells relative to luminal breast cancer cell lines (MCF7, T47D, HCC1500, CAMA1, EFM19, and ZR-75-1). Paired samples are connected with black lines. \**P* < 0.05 by paired one-sided *t* test after arcsine transformation of percentages.

To evaluate cell states more comprehensively, we profiled the transcriptomes of the six patients with paired tumor scrapes, zero-passage organoids, and 14-day 2D cultures (see Methods). Luminal breast cancer lines were included in the RNA sequencing for reference as before. Anticipating a mixture of cell lineages in the tumor scrapes, we used downsampled single-cell RNA-sequencing data from human breast cancers [21] to define a signature matrix for bulk deconvolution of 10 major cell types by CIBERSORTx [33] (Fig. 2C,D and File S2–S4). Tumor scrapes were predominantly luminal cells but contained detectable contributions from stromal cells, basal cells, endothelial cells, CD8+ T cells, macrophage–dendritic cells, and adipocytes (Fig. S1). When the same approach was applied to *ex vivo* cultures, we found that proliferative signatures were comparable to the tumor scrape but significantly less than established cell lines (Fig. S2). After 14 days, we found that zero-passage organoids generally retained the proportion of luminal cells, whereas 2D cultures did not (Fig. 2E). Both formats led to an increased percentage of basal cells, but the change was more exaggerated in 2D cultures compared to zero-passage organoids (Fig. 2F), illustrating the importance of matrigel-related cues for retaining luminal identity.

As a final characterization, we performed differential-expression analysis on the CIBERSORTx-inferred epithelial transcriptomes (File S5 and S6). In the analysis, basal and luminal lineages were combined because distinctions finer than epithelial–nonepithelial reduced the number of genes deconvolvable by CIBERSORTx. Focusing on the large gene expression changes (log2 fold change ≥ 2, log10 counts per million ≥1), we found 51 transcripts increased in zero-passage organoids compared to tumor scrapes and 410 transcripts decreased (FDR-corrected *P* < 0.05). Among the increases, we noted genes involved in Wnt–β-catenin regulation (*FRZB, SFRP1*), which likely related to feedback from Wnt3a in the Type 2 organoid medium [12]. There was also extensive upregulation of transcripts related to cholesterol metabolism (*DHCR7*, *EBP*, *MVD*, *NPC2*; File S7) and receptor tyrosine kinase signaling (*EFNB2*, *KIT*, *MAPK4*). Downregulated transcripts included many residual contaminants of non-epithelial genes incompletely resolved by CIBERSORTx (*HAVCR2*, *FCGR2B*, *PECAM1*, and others; File S7). However, decreases tied to X inactivation (*XIST*, *FTX*) and innate immunity (*TLR2*, *TLR4*) were likely cell autonomous and suggest a degree of reprogramming induced by the organoid format.

We extended the differential-expression analysis to compare zero-passage organoids with 14-day 2D cultures. Upon relaxing the magnitude of change to log2 fold change ≥ 1, we identified 29 upregulated and 46 downregulated transcripts. As expected, matrigel embedded organoids increased genes involved in polarity and membrane dynamics (*PARD6A*, *DOCK1*, *IQGAP2*), whereas 2D cultures on tissue-culture plastic increased genes for adhesion and F-actin stabilization (*ITGB1*, *LIMA1*, *COTL1*). More surprising was the upregulation of transcripts for proteins secreted by luminal epithelial cells in zero-passage organoids (*NUCB2*, *SCGB2A1*). 2D cultures, by contrast, were increased for various atypical MAPK genes (*MAP4K4*, *MAP3K20*, *MAPK6*) warranting further study. A complete set of differential-expression analyses is provided in File S6 along with gene set enrichment analyses in File S7.

### Clinical predictors of zero-passage organoid culture success

We assembled a cohort of 90 luminal breast cancers (Table 1) to determine if any pre-surgical characteristics predicted organoid culture success. Proportional hazard modeling detected enhanced success from cases with higher-grade tumor biopsies (*P* < 0.001; Fig. 3) and surgical resections (*P* = 0.045; Fig. S3). Cases obtained by mastectomy were also more successful compared to lumpectomy specimens (*P* ≤ 0.002; Fig. 3 and Fig. S3), which is counterintuitive given that mastectomy surgery times are often longer. Surgery-associated ischemia may trigger adaptive stress pathways that allow tumor cells to withstand the microenvironment change of organoid culture. We detected no impact of age, race, carcinoma type, hormone receptor expression, estimated tumor size, surgeon, pathologist, or culturist on organoid success (Fig. 3 and Fig. S4). These results help inform power analyses of future studies that apply zero-passage organoids to specific luminal cohorts.

**Table 1.** Patient demographic and clinical characteristics.

|  |  |  |  |  |  |  |  |  |
| --- | --- | --- | --- | --- | --- | --- | --- | --- |
| **Demographics** | **Sex** |  | **Race** |  | **Age** |  | **Presurgery Tx** |  |
|  | Female | 88 | White | 75 | ≤50 | 22 | None | 84 |
|  | Male | 2 | Black | 10 | 51–65 | 31 | Endocrine | 6 |
|  |  |  | Other | 5 | >65 | 37 |  |  |
| **Markers** | **ESR1<sup>†</sup>** |  | **PGR** |  | **HER2** |  |  |  |
|  | 0% | 1 | 0% | 6 | Negative | 87 |  |  |
|  | 50–95% | 41 | 0–50% | 28 | Positive | 3 |  |  |
|  | >95% | 48 | >50% | 58 |  |  |  |  |
| **Diagnostic core biopsy** | **Histologic Dx** |  | **Grade** |  | **Estimated size by imaging** |  |  |  |
|  | IDC | 71 | 1 | 32 | <1 cm | 22 | 3 to <4 cm | 7 |
|  | ILC | 14 | 2 | 49 | 1 to <2 cm | 28 | 4 to <5 cm | 2 |
|  | Other | 5 | 3 | 6 | 2 to <3 cm | 23 | 5+ cm | 10 |
|  |  |  | N/A | 3 |  |  | N/A | 8 |
| **Surgical specimen** | **Histologic Dx** |  | **Grade** |  | **Measured pathologic size** |  |  |  |
|  | IDC | 68 | 1 | 22 | <1 cm | 18 | 3 to <4 cm | 6 |
|  | ILC | 17 | 2 | 56 | 1 to <2 cm | 34 | 4 to <5 cm | 3 |
|  | Other | 5 | 3 | 12 | 2 to <3 cm | 22 | 5+ cm | 7 |
|  | **Surgery** |  | **Tumor stage** |  | **Nodal status** |  |  |  |
|  | Lumpectomy | 66 | pT1 | 55 | pN0, pNX | 76 |  |  |
|  | Mastectomy | 24 | pT2 | 28 | pN1 | 13 |  |  |
|  |  |  | pT3 | 7 | pN2, pN3 | 1 |  |  |

<sup>†</sup>Abbreviations: ESR1, estrogen receptor; PGR, progesterone receptor; HER2, human epidermal growth factor receptor 2; IDC, invasive ductal carcinoma; ILC, invasive lobular carcinoma; N/A, not available.

**Figure 3.**
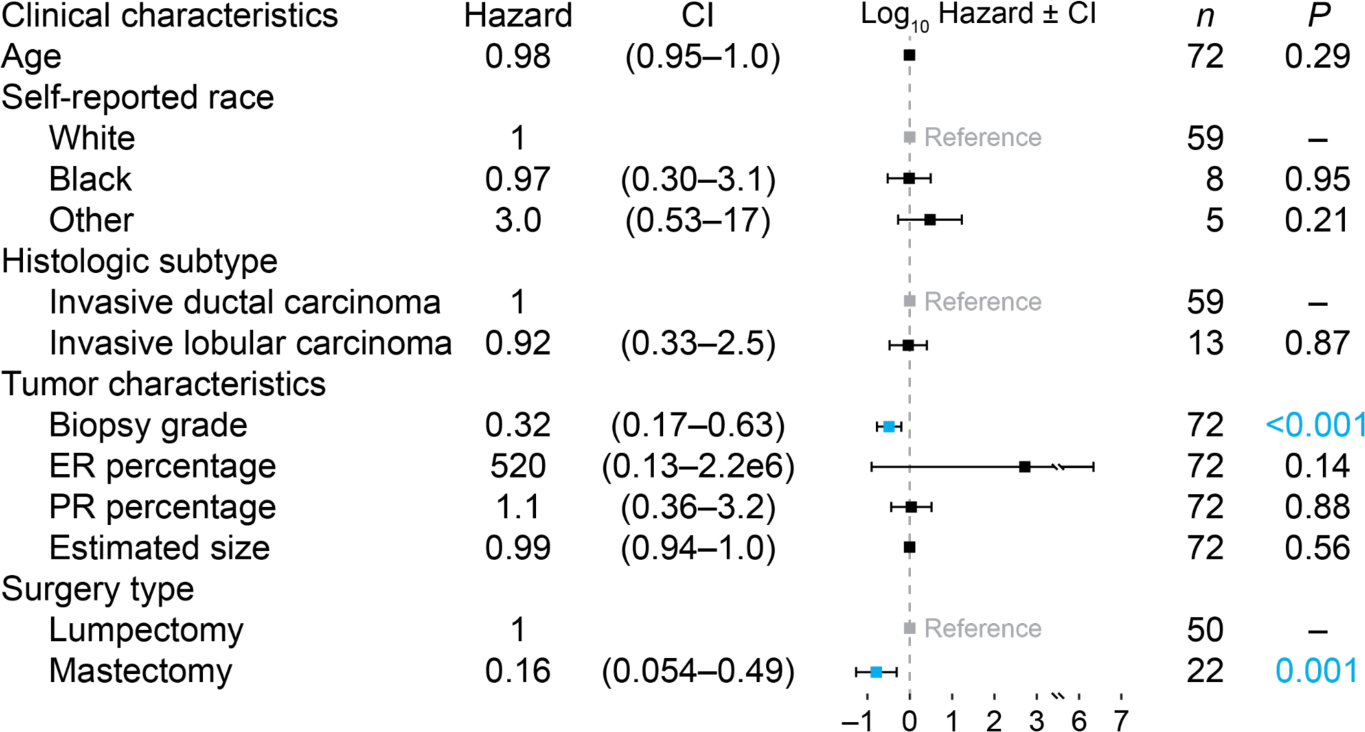
Pre-surgical characteristics and their association with organoid failure risk by day 14. Cox proportional hazards model for *n* = 72 luminal breast cancers showing hazard ratios with confidence interval (CI). Significant factors are blue. Tumor grade at biopsy was missing for one of the specimens, and five specimens were missing the estimated size (see Methods).

### Multiplexed zero-passage organoids by miniaturization

One tradeoff of the zero-passage approach is that all perturbations and controls must occur without appreciably expanding organoids from the tumor scrape. It was thus critical to shrink the culture volume as much as possible to allow multiple replicates and carefully assess the well-to-well variability of technical replicates. Prior studies did not describe culture volumes of matrigel less than 10–20 µl [12], prompting us to test reduced volumes of 7, 5, and 2 µl (*n* = 6 independent cases). We monitored growth of the organoid population longitudinally by serial brightfield imaging at six time points over 14 days, followed by digital image segmentation [16] and multiway ANOVA applied to the resulting distributions of organoid size (estimated by cross-sectional area). The ANOVA tracked fixed effects (patient, culture day, culture volume, and technical replicate), nesting (replicates within volume and volumes within patient), and pairwise interactions between factors [34]. Because post-hoc comparisons are challenging in multiway settings [35], we considered the volumes together and as individual pairs after correcting for multiple-hypothesis testing (Fig. 4A; see Methods). The analysis assessed differences by volume while accounting for covariates intrinsic to the study.

**Figure 4.**
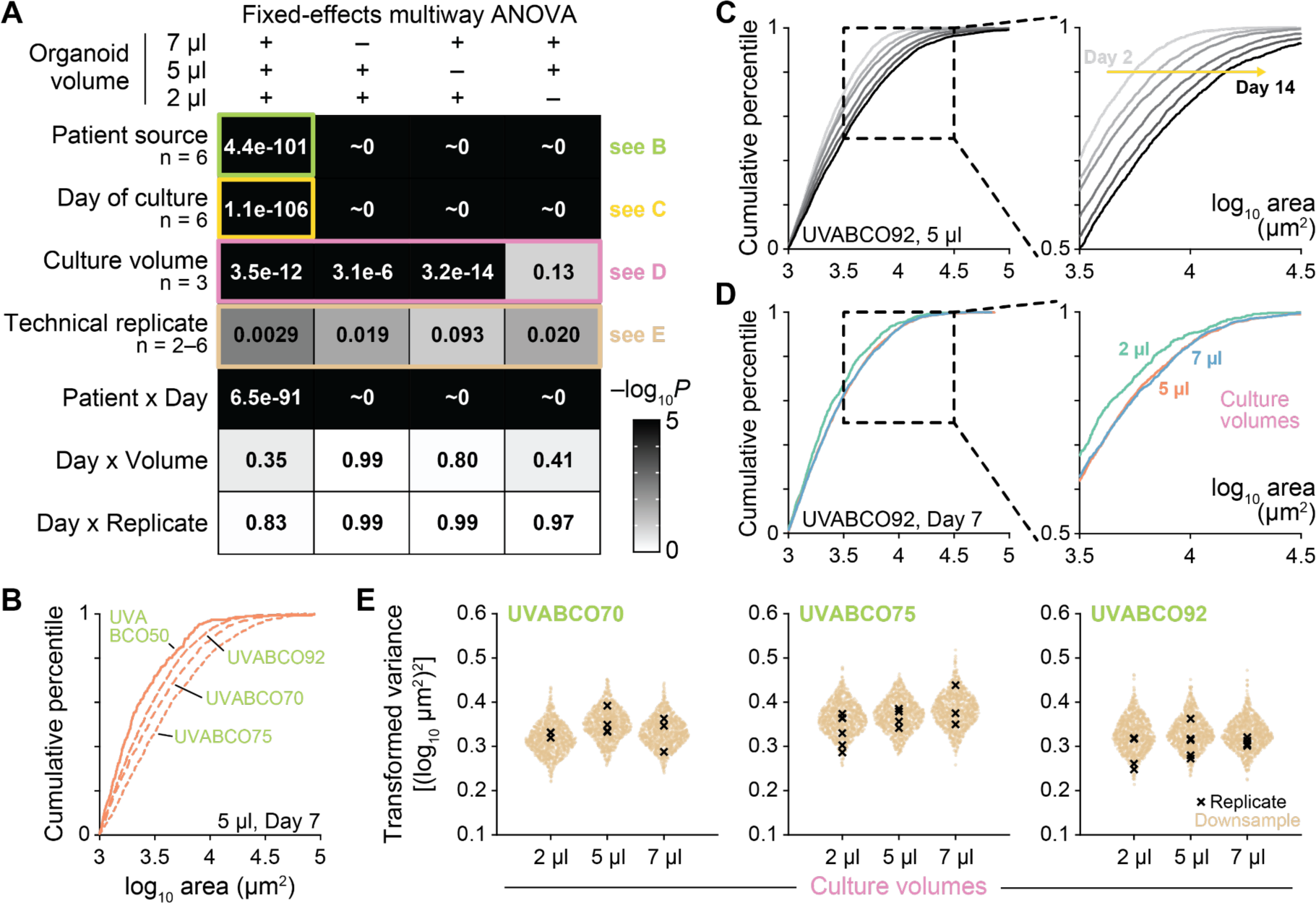
Reproducible miniaturization of organoid cultures to 5 µl without detectable growth deficits. **(A)** Multiway ANOVA table of *P* values for organoid size and the indicated fixed effects and interaction terms. Culture volumes (2 µl, 5 µl, and 7 µl) were considered together and in pairs (*n* = 2–6 factors or levels per effect). Specific illustrations of fixed effects are shown in subsequent subpanels. **(B)** Differences in organoid size distribution for four representative patient sources. 5 µl organoid cultures were measured for cross-sectional area by brightfield imaging on day 7. **(C)** Differences in organoid size distribution at 2, 5, 7, 10, 12 and 14 days. The 5 µl organoid culture was measured longitudinally as in (B). **(D)** Differences in organoid size distribution between 2 µl, 5 µl, and 7 µl cultures. The patient-derived sample was measured on day 7 as in (B). **(E)** Overlapping variance of downsampled technical replicates among organoid culture volumes. Organoid counts in the larger volumes of three patients were repeatedly downsampled to the 2 µl median (*n* = 1000 iterations; tan), and variance was estimated after shifted log transformation (see Methods). Variance estimates of the entire organoid count from each volume and technical replicate are overlaid (black). See Fig. S6 for analysis of additional patients. For (B–D), size distributions are summarized by their cumulative density function.

Among factors, patient, day of culture, and the patient x day interaction were highly significant, reflecting inter-tumor differences in growth and the increases in organoid size over two weeks (Fig. 4A– C). Culture volume was detected as a significant factor when all volumes were analyzed or when 2 µl was compared to 5 µl or 7 µl separately (Fig. 4A). We also observed suggestive increases in organoid death and attachment to tissue-culture plastic at the periphery of the matrigel droplet in 2 µl cultures (Fig. S5). The contact angle of a 2 µl culture is likely too small to prepare a hanging drop [12]. Notably, volume effects did not reach statistical significance when 5 µl and 7 µl cultures were compared, indicating negligible differences between these droplet sizes (Fig. 4D). For all culture volumes, variability among technical replicates was detectable but much less so compared to other factors (Fig. 4A). Volume-dependent replicate effects could not be estimated because of nesting. As a substitute, we downsampled organoid counts to the 2 µl median and inspected the distribution of 1000 iterations by patient, observing no consistent pattern with volume (Fig. 4E and Fig. S6). These results support miniaturization of organoid cultures down to 5 µl, doubling to quadrupling the number of replicates that can be generated from the limited material of a tumor scrape. Successful isolations yielded about 12 separate 5 µl drops (median; interquartile range: 6–29) for parallel testing.

### Zero-passage luminal breast cancer organoids exhibit patient-specific sensitivity to hormone interventions

One intended use of zero-passage organoids is for gauging the importance of hormone receptor signaling across different luminal breast cancers. Organoids are cultured in 100 nM β-estradiol [12] and the phenol red in the medium confers weak estrogen-like activity [36]. We thus designed a small panel of hormone interventions by withdrawing β-estradiol from the culture medium (to mimic aromatase inhibition that occurs when women take oral aromatase inhibitors in the adjuvant setting) and supplementing with progressive concentrations of 4-hydroxytamoxifen (4-HT) to block residual signaling. Before interventions, we cultured organoids for four days in 100 nM β-estradiol to disentangle growth from confounding effects on organoid initiation. The four-day preculture enabled formulation of a growth model linking cellular doublings to organoid cross-sectional areas based on an initial size at *t* = 0 days shared by all interventions (see Methods).

For six organoid preparations derived from five patients, we quantified cellular doubling times ranging from less than one week to over one month (Fig. 5A–F). Some cases appeared completely insensitive to tamoxifen (Fig. 5A,F) or only showed responses at the higher dose (Fig. 5B–D). We also identified instances of accelerated growth when β-estradiol was withdrawn (Fig. 5C–F), a possible reflection of the tumor “flares” reported clinically upon treatment with aromatase inhibitor [37, 38] or inhibitory effects of β-estradiol on breast cancer cell proliferation [39, 40]. In one patient with luminal breast cancers coming from each breast, we quantified a ∼twofold difference in growth rate between the tumors but the same pattern of responsiveness to β-estradiol withdrawal and tamoxifen (Fig. 5C,D). Importantly, the effects of β-estradiol removal and tamoxifen treatment were separable: UVABCO93 was affected by removal of β-estradiol but not tamoxifen (Fig. 5F), while the growth rate of UVABCO85 was altered only by higher dose of tamoxifen (Fig. 5B). These results indicate that the zero-passage timeline is sufficient to distinguish patient-specific differences in luminal cancer cell growth among alternative treatment regimens tested in parallel.

**Figure 5.**
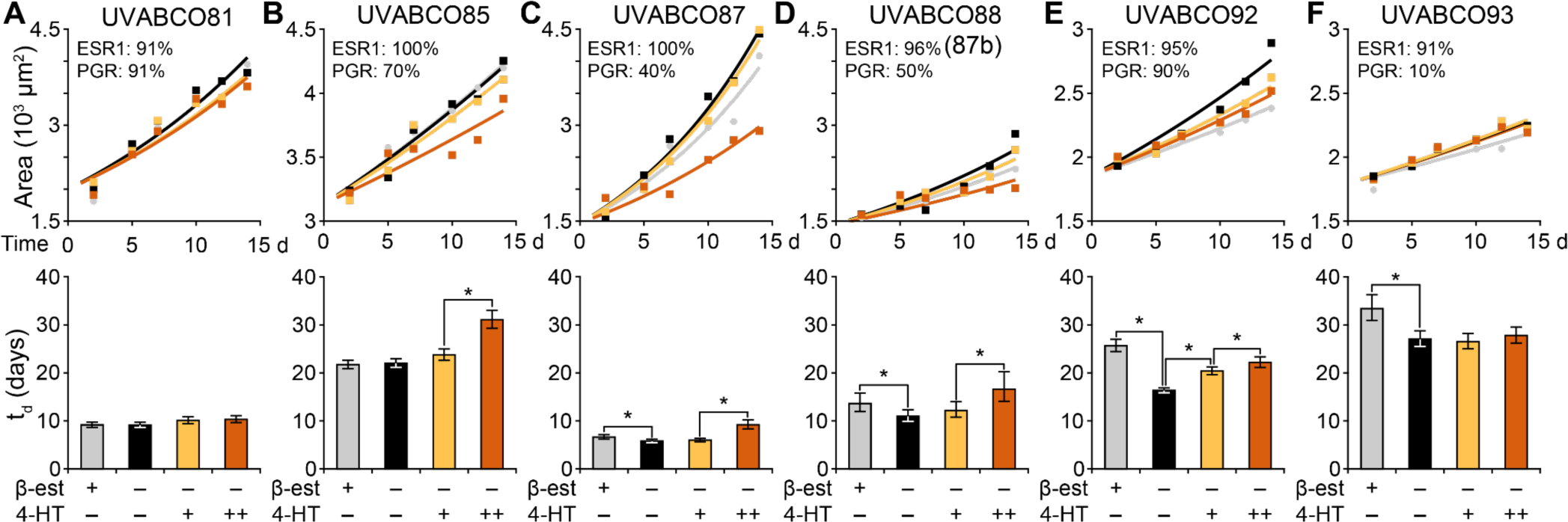
Patient-specific sensitivity of zero-passage organoids to hormone withdrawal and tamoxifen treatment. (**A**–**F**) Upper panels show mean organoid area at six time points after shifted log transformation (*n* = 22– 1407 organoids per patient, time point, and condition; see Methods) for control cultures (gray), cultures with β-estradiol withdrawn (β-est; black), and cultures with β-est withdrawn plus 200 nM 4-hydroxytamoxifen (4-HT; yellow) or 3 µM 4-HT (orange) on day 5. Overlaid are non-linear least-squares curve fits for area growth as a function of cellular doubling time (t_d_). Pathologic estimates of positivity for estrogen receptor (ESR1) and progesterone receptor (PGR) are reported as cellular percentages in the inset. Lower panels summarize the best-fit t_d_ for each condition with 87% confidence intervals estimated by support plane analysis. \**P* < 0.05 after Bonferroni correction for multiple-hypothesis testing.

### Highly efficient lentiviral transduction of luminal breast cancer organoids

To inform luminal cancer biology, zero-passage organoids must also be compatible with genetic manipulations. Luminal breast epithelial cells are difficult to transfect or transduce efficiently without additional steps (electroporation, enzyme treatments) that could stress freshly isolated cells [12, 41]. We achieved high transduction efficiency by resuspending pellets of standard lentiviral preparations in 1/10th volume of organoid growth medium, then incubating cells with virus for one hour before matrigel embedding and during the first three days of organoid culture (see Methods). Using this procedure with a mixture of EGFP- and TdTomato-encoding lentiviruses, we found that most organoids contained cells expressing one or more fluorophores (Fig. 6A,B). Importantly, transduction did not detectably affect organoid growth compared to paired untransduced controls or fluorescent-negative organoids within the same culture (*P* = 0.22; Fig. 6B). These results indicated that the lentiviral transduction protocol was gentle and effective enough to omit antibiotic selection [12], which if used would delay the zero-passage approach past two weeks.

**Figure 6.**
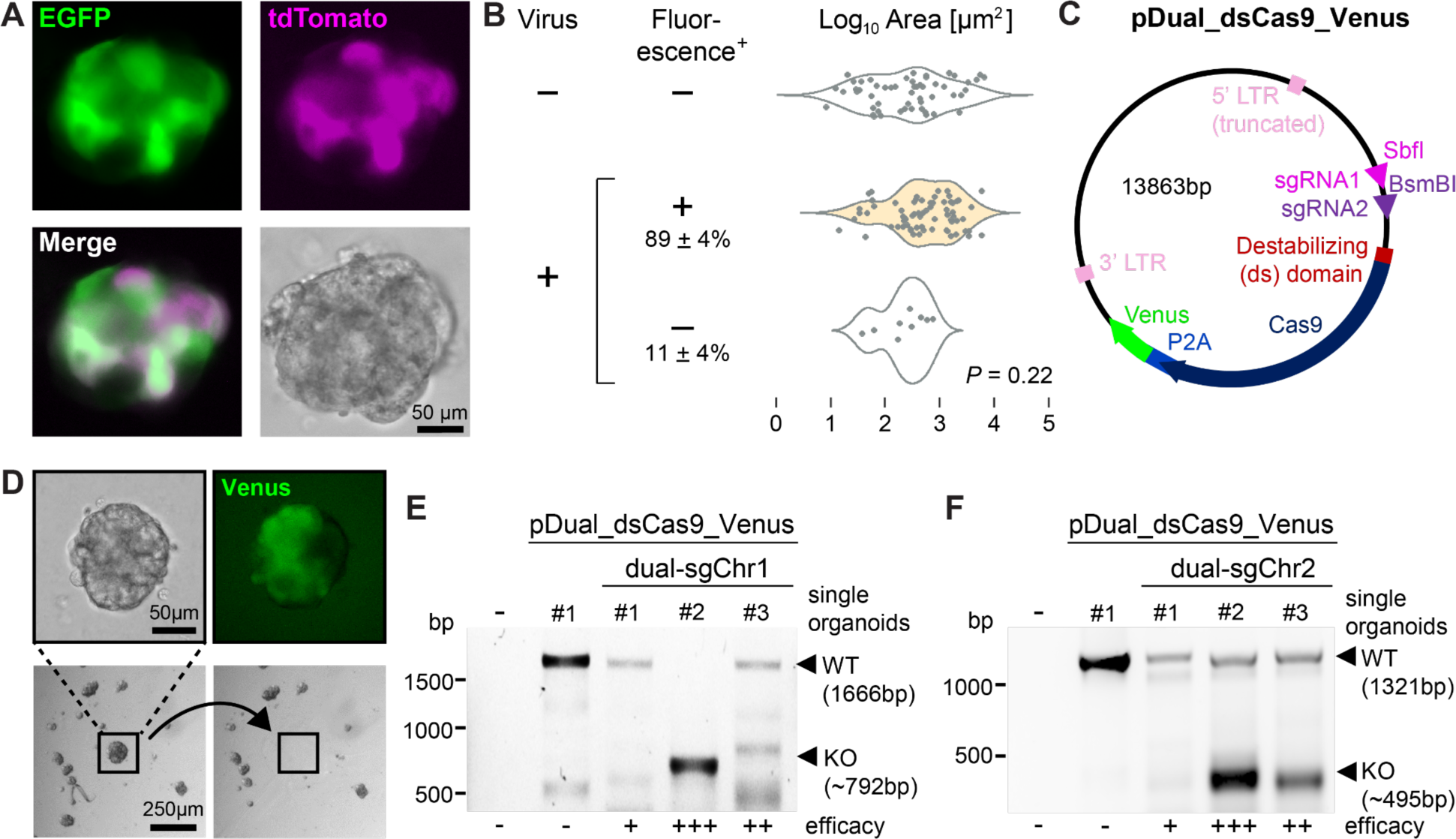
Zero-passage organoid compatibility with lentiviral transduction and genetic modification. **(A)** Representative image of one organoid transduced with a mixture of lentiviruses encoding EGFP (green) and TdTomato (magenta). Scale bar is 50 µm. **(B)** Quantification of organoids containing cells expressing one or more fluorophores. Percent fluorescence positive (+) is reported as the mean ± standard deviation from *n* = 4 independent luminal breast cancers. Differences in organoid sizes were assessed by two-way ANOVA (*n* = 11–27 organoids per group). **(C)** Map of the pDual_dsCas9_Venus lentiviral plasmid encoding destabilized Cas9 fused by a P2A sequence with Venus. Cloning sites for coexpressed dual sgRNAs are indicated. **(D)** Harvesting single Venus-positive organoids for genotyping. Fluorescent organoids confirmed by epifluorescence (top right) are harvested by micropipette aspiration (before: bottom left; after: bottom right). Scale bars are 50 µm (top) and 250 µm (bottom). (**E** and **F**) Single-organoid genotyping of zero-passage cultures transduced with pDual_dsCas9_Venus targeting a noncoding region of human chromosome 1 (**E**) or chromosome 2 (**F**). The first lane is a PCR negative control (no genomic DNA), the second lane is a nontargeting control (single organoid transduced with empty pDual_dsCas9_Venus), and the subsequent three lanes are single organoids transduced with dual sgRNAs. DNA markers are on the left and PCR band sizes for wild-type and knockout amplicons are on the right. Relative efficacy of targeting is indicated on the bottom of the gel.

Loss-of-function testing with CRISPR-Cas9 in organoids normally requires selection and clonal isolation of knockouts [12, 42, 43], which is problematic for retaining luminal identity (Fig. 1D,E). The sustained expression of Cas9 was also concerning as it activates the TP53 pathway and selects for inactivating mutations even without sgRNA-targeted double-strand breaks [44]. *TP53* mutations naturally occur in less than 30% of luminal breast cancers [28], raising the possibility of artifacts. We thus pursued a regulatable approach that modified an existing one-plasmid, destabilized Cas9-P2A-Venus system [45] to include a second sgRNA (Fig. 6C). Transient Cas9 stabilization should minimize off-target effects, and we reasoned that dual-sgRNAs—one targeting a critical exon and the other targeting a nearby intron—would result in more knockouts by the combined probability of indels, inverted reinsertions, and segmental deletions [46–48]. The P2A-linked reporter was also expanded by replacing Venus with mTagBFP2, puromycin resistance, or blasticidin resistance to create the pDual_dsCas9 series of lentivectors (see Methods). The collective modifications negligibly increased the final cloned sequence between the long terminal repeats (LTRs, Fig. 6C), and lentivirus of sufficiently high titer was produced when packaging by lipofection and concentrating as before (see Methods).

To control for the effect of double-strand breaks, we cloned dual-sgRNAs for non-coding regions of chromosome 1 or 2 that are considered safe targets [49] (see Methods). After transduction of zero-passage organoids and short-term Cas9 stabilization for 7 days, we detected Venus-positive organoids that were harvested by micropipette aspiration (Fig. 6D). Genotyping of single-organoid aspirates indicated a spectrum of segmental deletions that varied by organoid and dual-sgRNA control (Fig. 6E,F). Given that such genotyping overlooks small indels and inversions, we targeted a coding gene in luminal breast cancer lines as a proof of concept: the TP53 family member, *TP73*. The long variant of TP73 contains a transactivation (TA) domain with tumor-suppressive functions that can substitute for TP53 deficiency in breast cancer lines [50, 51]. *TP73* transcript abundance also exhibits high intratumor heterogeneity in many primary luminal breast cancers [52]. We designed dual-sgRNAs flanking the entire TA domain of TP73 (Fig. S7A) and transduced luminal breast cancer lines that were TP53 proficient (MCF7) or deficient (T47D). Non-concentrated pDual_dsCas9 lentivirus readily transduced cultured cells and achieved considerable *TP73* segmental deletions in selected (or sorted) pools of clones (Fig. S7B,C). To estimate the combined proportion of segmental deletions and smaller indels, we immunoblotted the polyclonal samples and quantified at least a twofold reduction in TP73 protein regardless of the P2A-linked reporter (Fig. S7D). We conclude that the pDual_dsCas9 lentivector series is an appropriate tool for genetically modifying organoids within a zero-passage time frame.

### Combinatorial phenotyping of genetic depletion and small-molecule treatment in zero-passage organoids

Zero-passage organoids can test mechanisms of action by combining genetic and small-molecule perturbations. As an illustration, we pursued one gene widely implicated in resistance to anti-cancer therapies: the NAD(P)H quinone dehydrogenase 1, *NQO1* [53, 54]. As part of the phase II antioxidant response, *NQO1* is heterogeneously upregulated in luminal breast cancers [52, 55]. The most potent chemical inducer of *NQO1* is sulforaphane (SFN), an electrophile and neutraceutical that has been tested as an antiproliferative agent for breast premalignancies (NCT00982319). SFN acts by inhibiting a specific ubiquitin ligase for the transcription factor, NFE2L2 [56–58]. SFN indirectly stabilizes NFE2L2 and inhibits proliferation of breast cancer cell lines in culture [59]. However, NQO1 promotes tumor growth *in vivo* through downstream effects on hypoxia sensing that are also relevant to large (>200 µm) organoids [60, 61]. Given that SFN-stabilized NFE2L2 upregulates many genes in addition to *NQO1* [62], we sought to ascertain the contribution of *NQO1* to SFN-induced responses in zero-passage organoids.

We disrupted *NQO1* by designing dual-sgRNAs flanking exon 2 and most of exon 3 upstream of its substrate-binding domain (Fig. 7A) and transducing seven independent luminal cases in parallel with safe-targeting controls. Cas9 was stabilized in zero-passage organoids for the first seven days, and different extents of segmental deletion were confirmed in single organoids (Fig. 6E,F and 7B). After seven days, half of the replicates were treated with SFN at a clinically-attainable concentration of 10 µM [63] and parallel cultures monitored for another seven days. Because viral transductions began at *t* = 0 days, we expanded the growth model to account for different starting points of each genotype and estimated doubling times globally (see Methods).

**Figure 7.**
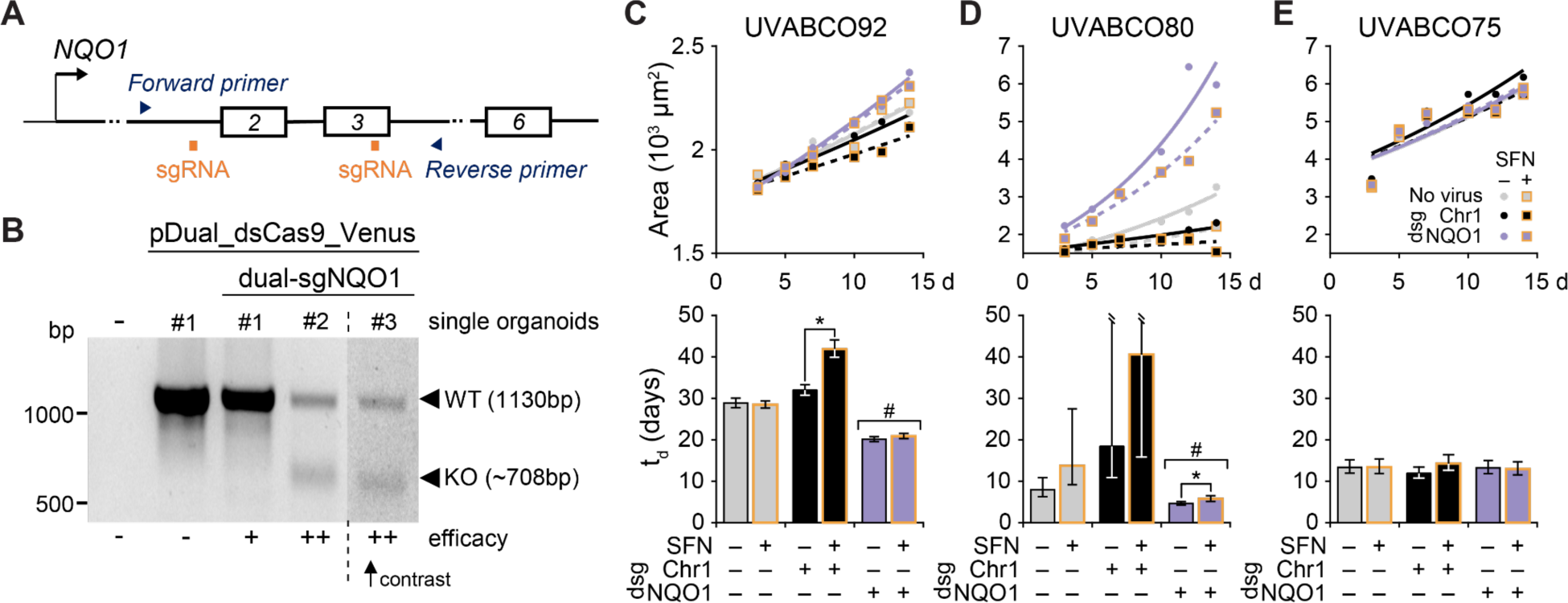
Combinatorial *NQO1* depletion and sulforaphane treatment of zero-passage organoids yields three classes of patient-specific responses. **(A)** Experiment design for targeting and detecting deletion of *NQO1*. Dual sgRNAs targeting required exons are in orange and genotyping PCR primers are in blue. **(B)** Single organoid genotyping of zero-passage cultures transduced with pDual_dsCas9_Venus targeting *NQO1* as in (A). The first lane is a PCR negative control (no genomic DNA), the second lane is a nontargeting control (single organoid transduced with empty pDual_dsCas9_Venus), and the subsequent three lanes are single organoids transduced with dual sgRNAs. DNA markers are on the left and PCR band sizes for wild-type and knockout amplicons are on the right. Relative efficacy of targeting is indicated on the bottom of the gel. The rightmost lane was contrast-enhanced independently for better visibility of the bands due to low genomic DNA content. (**C**–**E**) Upper panels show mean organoid area at six time points after shifted log transformation (*n* = 17– 1661 organoids per patient, time point, and condition; see Methods) for control cultures transduced with Cas9 only (gray), cultures with dual-sgRNAs (dsgRNAs) targeting a noncoding region of chromosome 1 (Chr1) targeted (black), and cultures with *NQO1* targeted with dsgRNA (purple). Genotypes were treated with 10 µM sulforaphane (SFN, yellow boxes) or 0.1% DMSO control. Overlaid are non-linear least-squares curve fits for growth as a function of cellular doubling time (t_d_) and genotype. Lower panels summarize the best-fit t_d_ for each condition with 91% confidence intervals estimated by support plane analysis. \**P* < 0.05 after Bonferroni correction for multiple-hypothesis testing for the indicated pairwise comparisons. ^#^*P* < 0.05 indicating both dsgNQO1 conditions are less than the corresponding dsgChr1 condition. See Fig. S8 for analysis of additional patients.

We noted three classes of responses to *NQO1* knockout ± SFN treatment in zero-passage organoids. For three of seven cases, SFN elicited growth inhibition that was reduced or lost upon *NQO1* knockout (Fig. 7C and Fig. S8A,B). In another three of seven cases, *NQO1* knockout was enough to accelerate organoids growth (Fig. 7D and Fig. S8C,D). One case was unaffected by any perturbation (Fig. 7E). Neither *NQO1* nor *NFE2L2* are recurrently mutated in breast cancer, but they drive other cancers [64] and their non-oncogenic regulation [65] may confer patient-specific growth and adaptive properties. Combining zero-passage organoids with genetic modifications enables these properties to be rapidly interrogated.

## Discussion

Through speed and miniaturization, zero-passage organoids preserve luminal breast cancers long enough to distinguish patient-specific growth phenotypes after chemical or genetic perturbation. By hewing closely to standard clinical procedures, we successfully engaged a composite team of surgeons and pathologists without incurring detectable batch effects in organoid success. The scrape-to-organoid pipeline should be extendable to any medical center where surgery, pathology, and research facilities are nearby. The two-week time frame of zero-passage organoids aligns well with adjuvant decisions about therapy that do not occur until one month after surgery or later. In the future, it may be possible to inform early therapy decisions by iterating different treatment scenarios through zero-passage organoids in parallel. Immediately, the approach provides a reliable source of primary material to complement research in luminal breast cancer cell lines and animal xenografts.

Central to the zero-passage approach is parallelization through miniaturized replicates that do not sacrifice growth, viability, or technical reproducibility. The number of replicates and thus independent variables is limited by the total material from the tumor scrape and the fraction of that material per replicate. The lower limit of 5 µl might not be surmounted with smaller microwell plates—2 µl of matrigel in a low-volume 384-well plate creates a cylindrical plug no taller than the diameter of an average organoid. Smaller samples may also become less reflective of the overall tumor heterogeneity; we occasionally noted instances where culture volumes differed in organoid size variability (Fig. S2C). At the existing scale, we believe there is great potential to collect more readouts from zero-passage organoids. We used serial brightfield imaging and transcriptomics here but foresee alternative endpoints quantifying cell death, cytokine release, and protein localization. Clearly, organoid phenotypes should be focused on responses to the interventions planned.

A persistent challenge in working with luminal breast cancer cells *ex vivo* is loss of hormone receptors, which transcriptionally downregulate after just a few days of standard tissue culture [66]. Luminal breast cancer lines are de-differentiation outliers and represent an oversimplification of primary tumor states [67]. Organoid conditions do not fix the problem of de-differentiation [12, 31] but may postpone it long enough for zero-passage cultures to yield biological insight. Indeed, the tamoxifen responsiveness we often observed suggests that ESR1 signaling remains intact. Our paired comparison with monolayer-cultured cells indicates that decreases in hormone receptors are less severe when embedded in the adhesive and mechanosensory environment of matrigel. Traditional 2D culture and the unlimited access to oxygen, nutrients, metabolites, and signaling molecules it provides may accelerate hormone receptor decline by fostering proliferation over cell-cell and cell-matrix interactions typical of solid tumors [68, 69]. Looking ahead, it may be possible to extend the zero-passage window beyond two weeks by using alternative culture conditions [30, 70] or hydrogel matrices [71] designed to maintain hormone receptors.

## Conclusions

Zero-passage organoids consider each set of molecular and genetic perturbations as an n-of-1 study specific to a patient. In contrast to n-of-1 clinical trials, organoids accommodate paired designs with control, alternative, and interactive arms that strengthen local interpretations. Organoid phenotypes can support clinical decision-making without predicting long-term patient outcomes directly. Variation between zero-passage organoids derived from different patients is considerable, yet rigorous findings that are truly translatable should replicate nonetheless, with outliers probed more deeply by sequencing. Our study minimizes barriers to accessing authentic luminal breast cancer cells, examining their properties, and testing their vulnerabilities.

## Data availability

The plasmids used in the study are available at Addgene (see Methods). RNA-seq data files are available under GEO accession number GSE262110.

## List of abbreviations

2D: monolayer-cultured cells
4-HT: 4-hydroxytamoxifen
adDMEM+: Advanced DMEM/F12 supplemented with HEPES and GlutaMAX^TM^
β-est: β-estradiol
Blast: blasticidin resistance
CI: confidence interval
D-BSA: DMEM GlutaMAX containing penicillin–streptomycin and fatty acid-free BSA
DCs: dendritic cells
dsgChr1: dsgRNA targeting a noncoding region of human chromosome 1
dsgRNA: dual-sgRNA
dsgTP73: dsgRNA targeting *TP73*
GEP: gene expression profile
LTR: long terminal repeat
MΦs: macrophages
O: organoid culturist
P: pathologist
Puro: puromycin resistance
RT-qPCR: reverse transcription-quantitative polymerase chain reaction
S: surgeon
SFN: sulforaphane
sgRNA: single guide RNA
t_50_: median survival time
t_d_: cellular doubling time

## Acknowledgements

We are grateful to the collaborating surgeons (David Brenin, Lynn Dengel, Anneke Schroen) and pathology staff (Sarah Bastarache, William Lee Carstens, Lisa Friedman, Tappy Gish, Akriti Gupta, AnnaVi Jones, Lauren Luketic, Nicole McGinn, Margaret Moore, Ansley Scott, Selveras Zayed) who helped procure clinical material for this study. We thank Hui Zong, Amy Bouton, and Kenley Ellis for reviewing this manuscript, Jeffrey Hsu and Mariah Snelson for early assistance with the work, Wisam Fares and Andrew Sweatt for coding help, and Emily Farber and Suna Onengut-Gumuscu for performing RNA sequencing at the UVA Center for Public Health Genomics. Data were partly generated in the University of Virginia Flow Cytometry Core Facility (RRid:SCR_017829) that is partially supported by the UVA Comprehensive Cancer Center support grant (P30-CA044579). We acknowledge Research Computing at The University of Virginia for providing computational resources and technical support that have contributed to the results reported within this publication.

## Author information

### Authors and Affiliations

Department of Biomedical Engineering, University of Virginia School of Engineering, Charlottesville, VA 22908, USA

Róża K Przanowska, Najwa Labban, Piotr Przanowski, Russell B Hawes, Kevin A Janes

Department of Pathology, University of Virginia, Charlottesville, VA 22908, USA Kristen A Atkins

Department of Surgery, University of Virginia Health System, Charlottesville, VA 22908, USA Shayna L Showalter

UVA Comprehensive Cancer Center, University of Virginia, Charlottesville, VA 22908, USA. Shayna L Showalter, Kevin A Janes

Department of Biochemistry and Molecular Genetics, University of Virginia School of Medicine, Charlottesville, VA 22908, USA

Kevin A Janes

### Contributions

Conceptualization: KAJ, SLS, RKP, KAA, NL; Data curation: KAJ, RKP, NL, PP; Formal Analysis: KAJ, RKP, NL, PP; Funding acquisition: KAJ, SLS, RKP, NL, RBH; Investigation: RKP, NL, PP, RBH; Methodology: RKP, NL, KAJ; Project administration: KAJ, SLS; Resources: KAJ, SLS, NL, RKP, RBH; Supervision: KAJ, SLS; Validation: RKP, NL; Visualization: KAJ, RKP, PP, NL; Writing – original draft: RKP, KAJ, NL, PP; Writing – review & editing: SLS, KAA, RBH.

### Corresponding authors

Shayna L Showalter, Kevin A Janes

## Ethics declarations

### Ethics approval and consent to participate

Human sample acquisition and experimental procedures were carried out in compliance with regulations and protocols approved by the Institutional Review Board for Health Sciences Research (IRB-HSR) at the University of Virginia in accordance with the U.S. Common Rule and IRB Protocol #14176. The Institutional Review Board has granted this study a waiver of consent under 45CFR46.116 of the 2018 Common Rule.

### Consent for publication

Not applicable.

## Supplementary Information

**Figure S1.**
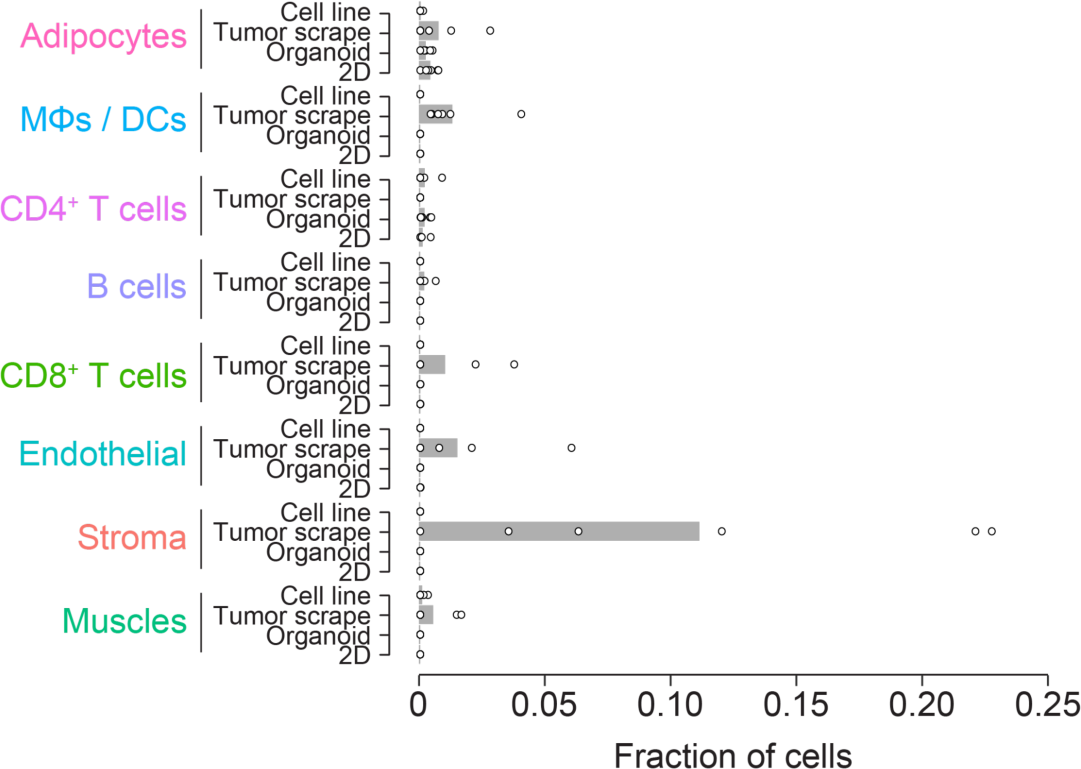
Deconvolution identifies non-epithelial cell types predominantly in tumor scrapes. CIBERSORTx-inferred cell lineage fractions (excluding Basal and Luminal shown in Fig. 2E,F) in originating tumor scrapes, zero-passage organoids, 2D cultured cells and luminal breast cancer cell lines (*n* = 6 samples in each group).

**Figure S2.**
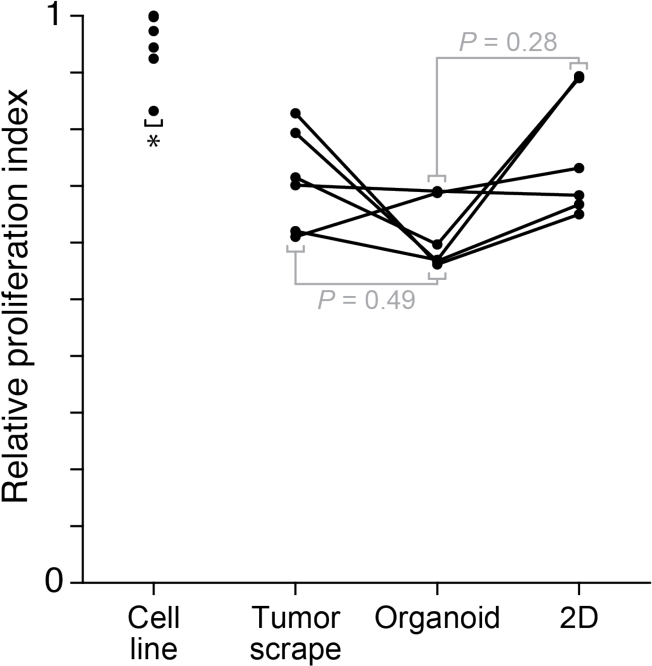
Luminal cell lines are more proliferative than tumor scrapes, zero-passage organoids, and short-term 2D cultures. The CIBERSORTx-inferred epithelial transcriptomes were projected on a gene signature for proliferation rate [23] and normalized to the maximum-observed rate of luminal breast cancer cell lines (MCF7, T47D, HCC1500, CAMA1, EFM19, and ZR-75-1). \**P* < 0.05 by paired two-sided *t* test after arcsine transformation of percentages and Šidák correction for multiple hypothesis testing.

**Figure S3.**
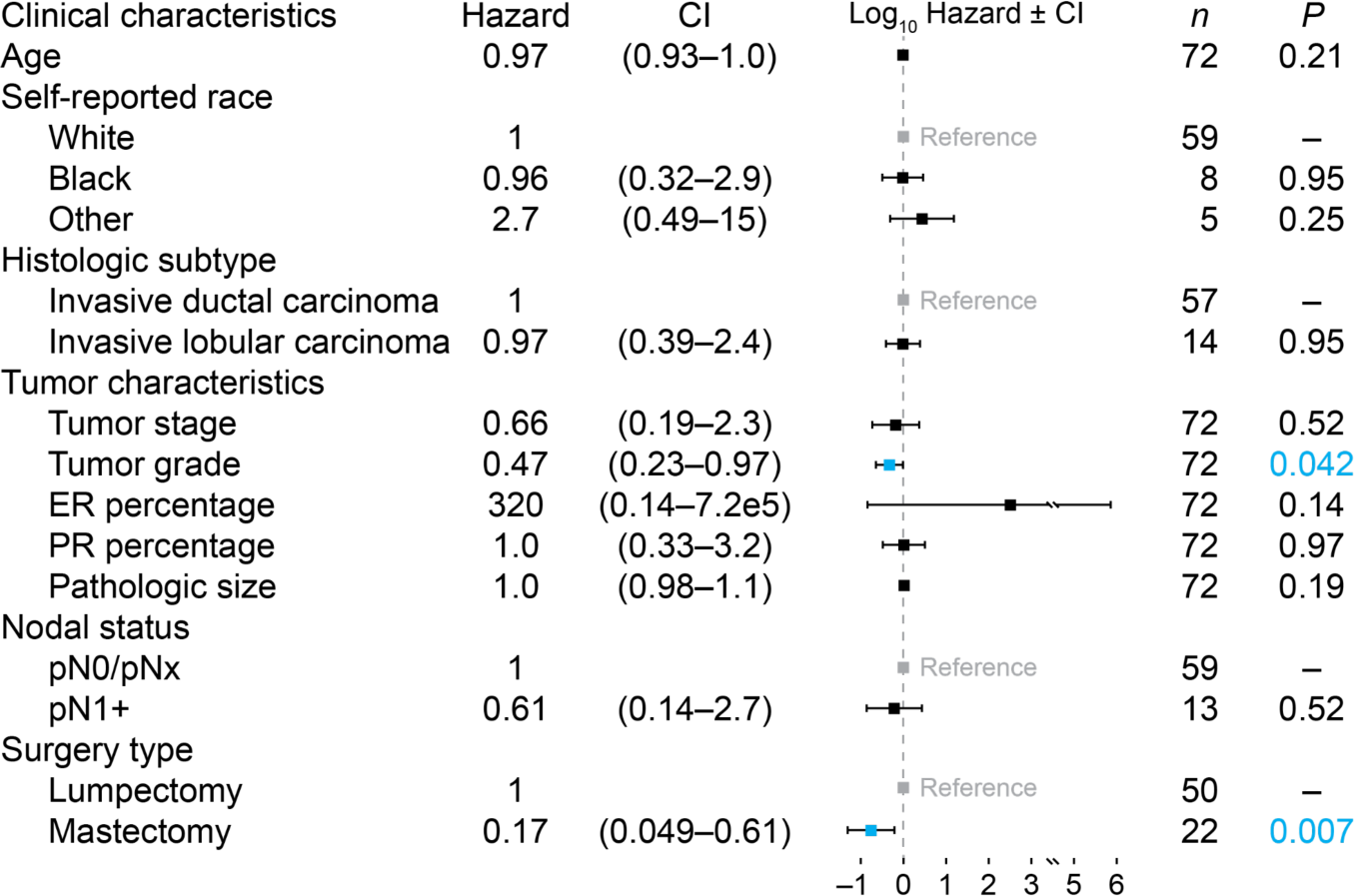
Post-surgical characteristics and their association with organoid failure risk by day 14. Cox proportional hazards model for *n* = 72 luminal breast cancers showing hazard ratios with confidence interval (CI). Significant factors are blue. One surgical case contained both intraductal carcinoma and intralobular carcinoma, which led to the exclusion of that factor for this case. Tumor grade at resection was missing for one of the specimens (see Methods).

**Figure S4.**
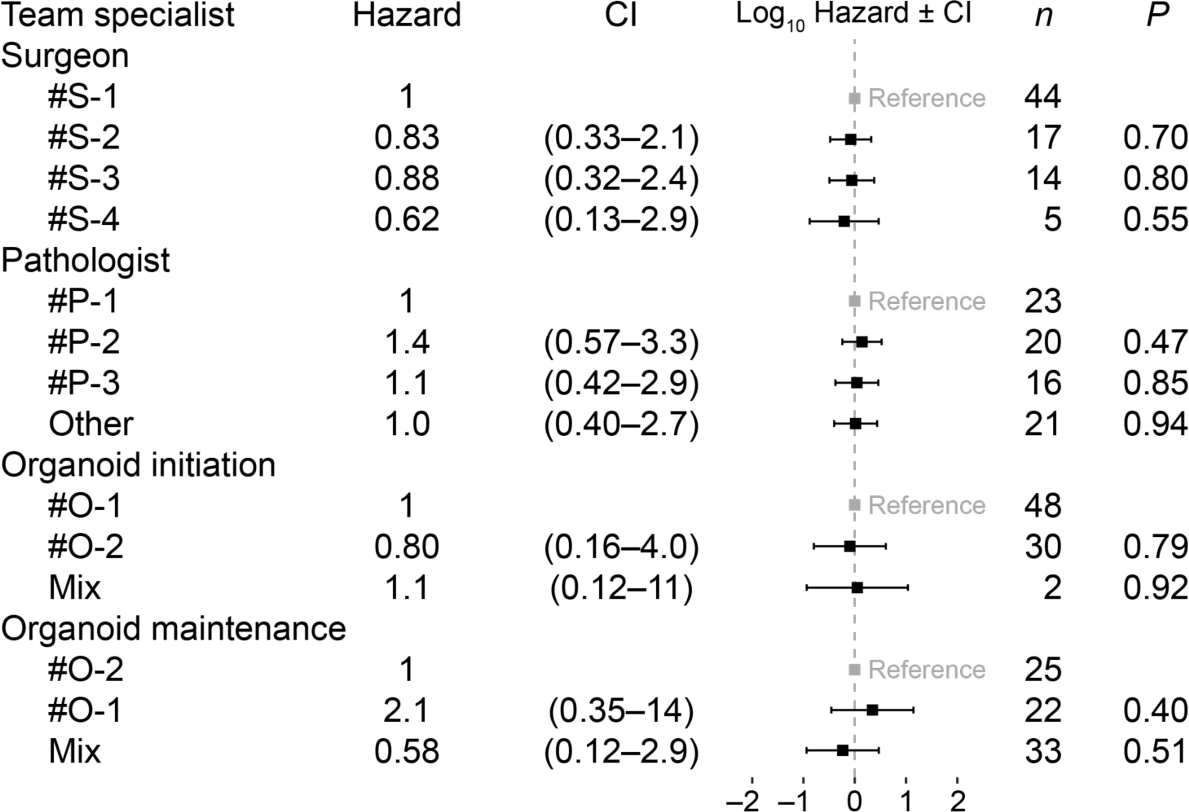
Handler characteristics and their association with organoid failure risk by day 14. Cox proportional hazards model for *n* = 72 luminal breast cancers showing hazard ratios for surgeon (S), pathologist (P), and organoid culturist (O) with confidence interval (CI). “Other” indicates 11 different pathology assistants. “Mix” indicates O-1 and O-2 worked on organoid initiation or maintenance together.

**Figure S5.**
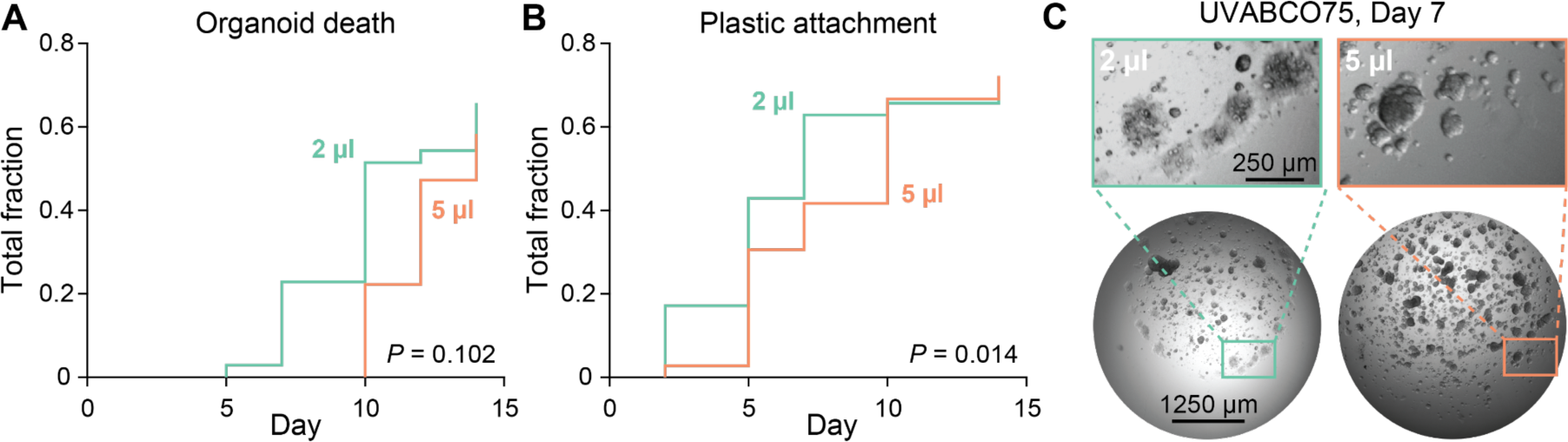
2 µl organoid cultures attach to plastic earlier and may die faster. (**A** and **B**) Cox proportional hazard models for organoid death and plastic attachment using volume as a continuous variable for *n* = 35–36 cultures from 12 independent luminal breast cancers. **(C)** Representative image of a paired 2 µl and 5 µl organoid culture with premature plastic attachment on day 7. Scale bars are 250 µm (top) and 1250 µm (bottom).

**Figure S6.**
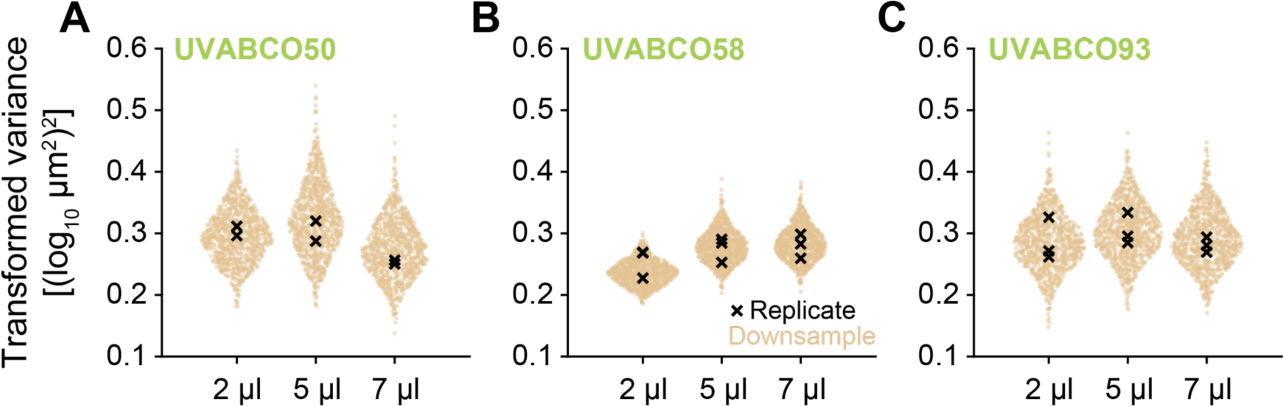
Differences in organoid size variance are not consistently attributable to culture volume. (**A**–**C**) Organoid counts in the larger volumes of three patients were repeatedly downsampled to the 2 µl median (*n* = 1000 iterations; tan), and variance was estimated after shifted log transformation (see Methods). Variance estimates of the entire organoid count from each volume and technical replicate are overlaid (black).

**Figure S7.**
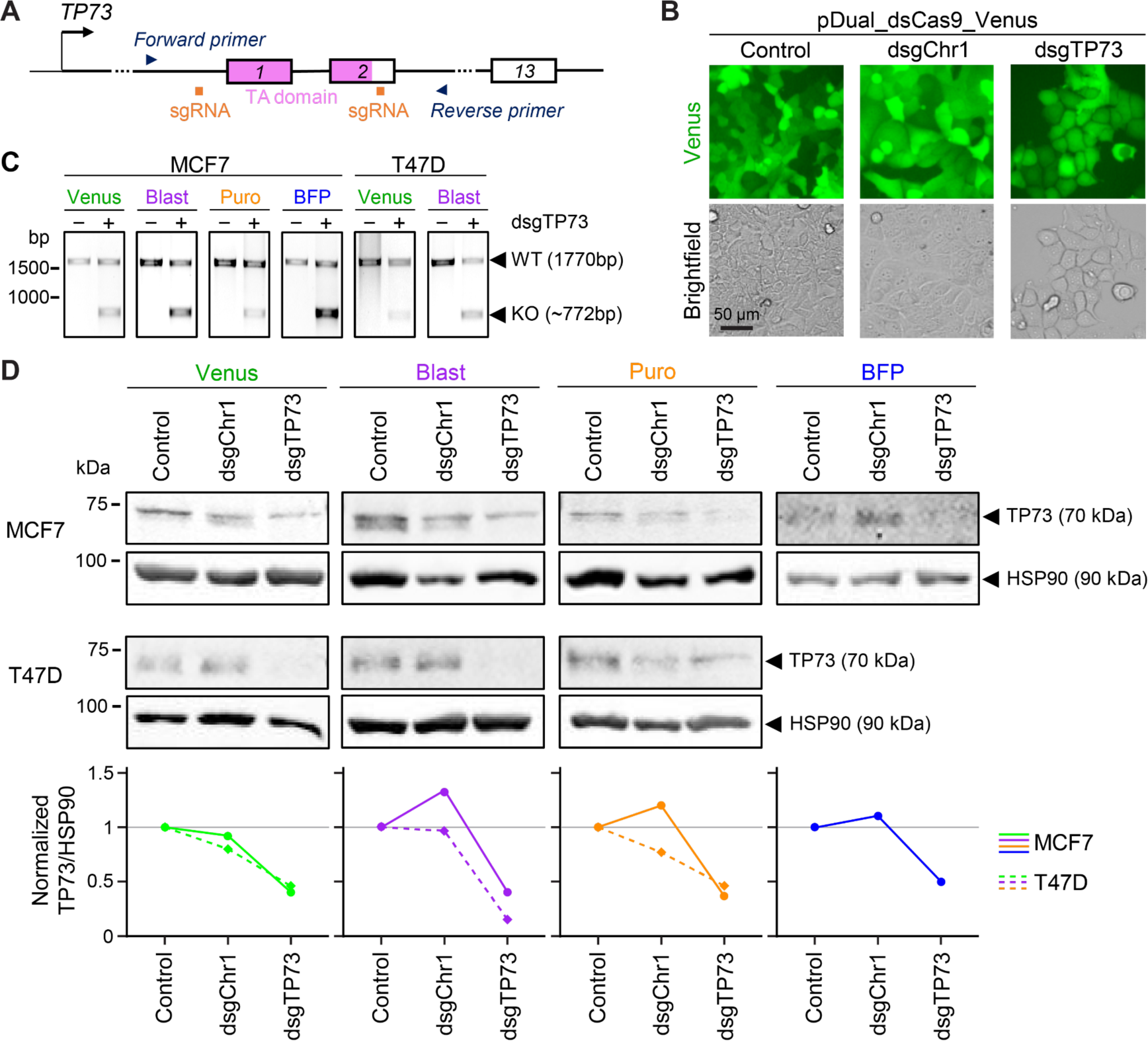
Efficient transduction of pDual_dsCas9_Venus and deletion of *TP73* in luminal breast cancer cell lines. **(A)** Experimental design for targeting and detecting deletion of the transactivation domain of *TP73* (TA, pink). Dual sgRNAs are in orange and genotyping PCR primers are in blue. **(B)** Confirmation of Venus-sorted MCF7 cells transduced with different versions of pDual_dsCas9_Venus. Scale bar is 50 µm. **(C)** Genotyping of selected polyclonal MCF7 or T47D cells transduced with different pDual_dsCas9 versions encoding dual-sgRNA (dsgRNA) of *TP73* as in (A). The control lane is empty pDual_dsCas9_Venus (nontargeting), and the second lane is from cells transduced and selected by sorting or antibiotic. DNA markers are on the left and PCR band sizes for wild-type and knockout amplicons are on the right. **(D)** Immunoblotting of TP73 and HSP90 (housekeeping control) in selected polyclonal MCF7 or T47D cells transduced with empty pDual_dsCas9 (control), pDual_dsCas9 encoding dsgRNA targeting a noncoding region of human chromosome 1 (dsgChr1), or pDual_dsCas9 encoding dsgRNA targeting *TP73* as in (A) (dsgTP73) sorted (Venus and BFP) or selected (Blast and Puro). Protein markers are on the left and proteins are on the right. Blots were quantified, normalized to HSP90, and plotted relative to control (gray line).

**Figure S8.**
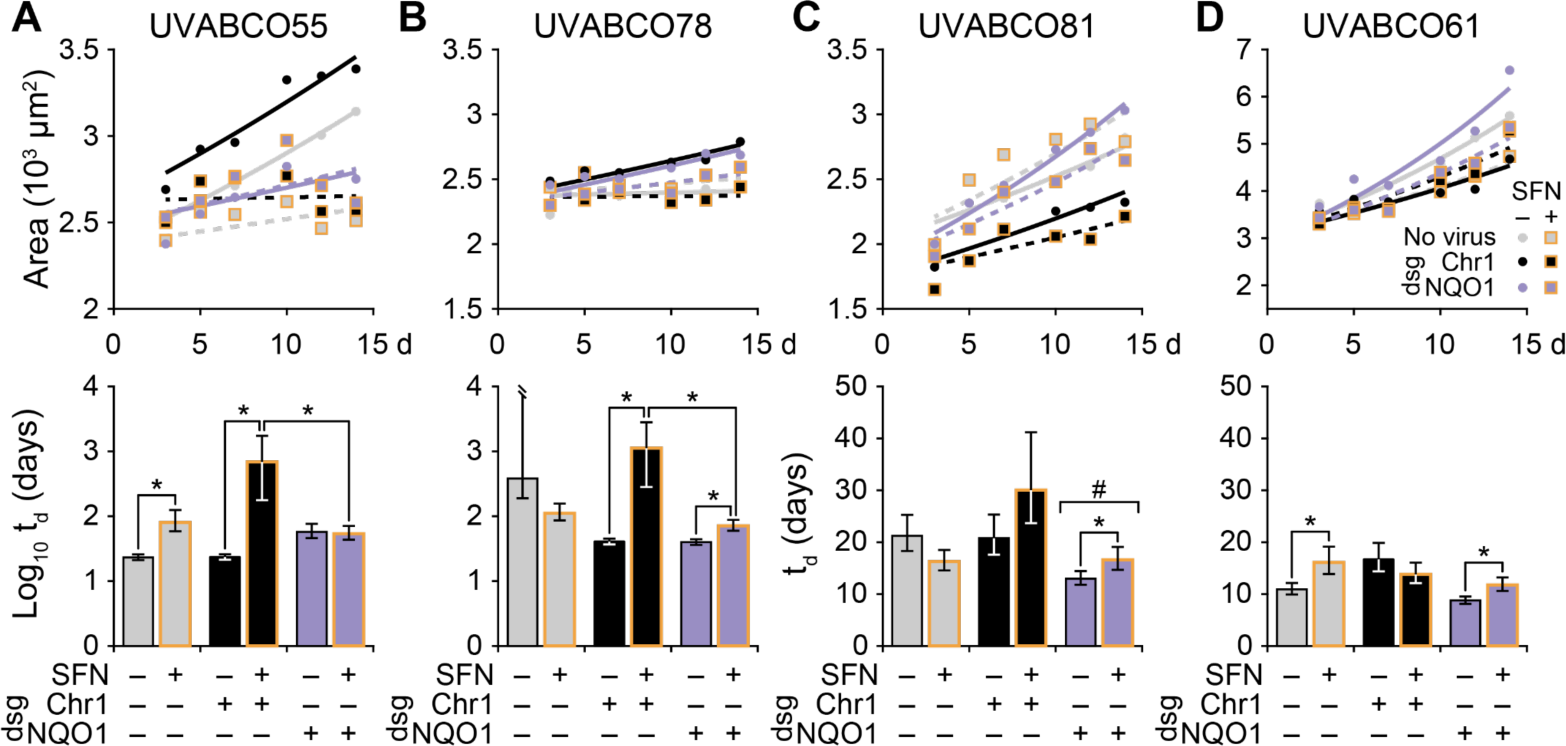
Different classes of patient-specific responses to combinatorial *NQO1* depletion and sulforaphane treatment in zero-passage organoids. **(A)** Upper panels show mean organoid area at six time points after shifted log transformation (*n* = 113– 1648 organoids per patient, time point, and condition; see Methods) for control cultures transduced with Cas9 only (gray), cultures with dual-sgRNAs (dsgRNAs) targeting a noncoding region of chromosome 1 (Chr1) targeted (black), and cultures with *NQO1* targeted with dsgRNA (purple). Genotypes were treated with 10 µM sulforaphane (SFN, yellow boxes) or 0.1% DMSO control. Overlaid are non-linear least-squares curve fits for growth as a function of cellular doubling time (t_d_) and genotype. Lower panels summarize the best-fit t_d_ for each condition with 91% confidence intervals estimated by support plane analysis. \**P* < 0.05 after Bonferroni correction for multiple-hypothesis testing for the indicated pairwise comparisons. ^#^*P* < 0.05 indicating both dsgNQO1 conditions are less than the corresponding dsgChr1 condition.

